# Multimodal transcriptomics and calcium imaging reveal a novel subset of polymodal nociceptors expressing the interleukin 1 receptor in mice

**DOI:** 10.64898/2026.07.21.739828

**Authors:** Camille Illiano, Dominic Bélanger, Harrison J. Stratton, Nicolas Vallières, Nadia Fortin, Martine Lessard, Louison Brochoire, Marine Christin, Yves De Koninck, Reza Sharif-Naeini, Sarah E. Ross, Manon Defaye, Feng Wang, Steve Lacroix

**Author notes:** Correspondence to: Dr. Steve Lacroix, Centre de recherche du CHU de Québec–Université Laval, 2705, boul. Laurier, Québec, QC, Canada, G1V 4G2.

## Abstract

We previously established that sensory neurons in the mouse dorsal root ganglia (DRG) express a functional receptor for the proinflammatory cytokine interleukin (IL)-1. We also demonstrated that deletion of the IL-1 receptor type 1 gene, *Il1r1*, in TRPV1-expressing (+) neurons prevented pain in models of chronic inflammatory diseases such as multiple sclerosis, rheumatoid arthritis and osteoarthritis. Here, we found a marked sex difference in the abundance of IL-1R1^+^ neurons, which represented approximately 10% of all DRG neurons in females but only 5% in males. However, male mice exhibited stronger, longer-lasting mechanical hypersensitivity than females after IL-1β injection into the cerebrospinal fluid. *In vivo* calcium imaging revealed that IL-1β- responsive DRG neurons responded to cutaneous mechanical and capsaicin stimulation. By integrating spatial transcriptomics with single-cell RNA sequencing (scRNA-Seq), we identified a gene signature uniquely marking IL-1R1^+^ neurons, including genes such as *Ada*, *Cysltr2*, *Gpr139*, *Htr1a*, *Htr1f*, *Il31ra*, *Nppb*, *Npy2r*, *Nts*, *P2rx2*, *Pde4c*, and *Sst*, with *Ada* and *Sst* validated at the protein level. Omics analysis revealed that IL-1R1^+^ DRG neurons form a subset of non-peptidergic type 3 (NP3) sensory neurons, which are linked to inflammatory pain and itch. However, *Il1r1* deletion did not affect itch responses to serotonin, histamine, or chloroquine, and these mediators failed to induce calcium activity in IL-1β-responsive DRG neurons. Finally, scRNA-seq identified several genes upregulated in NP3 neurons after IL-1β injection, including *Alkal2*, *Bdnf*, and *Lcn2*, associated with chronic inflammatory pain. Thus, our study unveils novel markers for IL-1R1^+^ nociceptors and reaffirms their selective role in inflammatory pain.

## INTRODUCTION

Chronic pain affects ∼20% of adults worldwide and is a leading cause of medical consultations (Goldberg and McGee, 2011). It is especially prevalent in people suffering from chronic inflammatory conditions, including neurodegenerative diseases and neurotrauma, where neuroinflammation is a key feature. For example, chronic pain is present in ∼50% of patients with multiple sclerosis (Mifflin and Kerr, 2017; O’Connor et al., 2008), while it affects 50-75% of individuals with traumatic brain injury, spinal cord injury (SCI), or peripheral nerve injury (PNI) (Gaudet et al., 2021; Miclescu et al., 2019; Nampiaparampil, 2008). Pain is transmitted through a specialized subset of sensory neurons called nociceptors, which have their cell bodies in dorsal root ganglia (DRGs) and send nociceptive signals via afferent nerve fibers from the periphery to the spinal cord and brain. Importantly, pain is a key symptom of inflammation, suggesting a causal link between them. Understanding how cells promoting inflammation communicate with nociceptors, and vice-versa, is therefore fundamental to comprehend and treat pain.

Nociceptors consist of a subset of neurons expressing specialized receptors that respond to mechanical, thermal and/or chemical stimuli. Pain perception arises from action potential initiation within these nociceptors. Perhaps the most studied of these receptors is the transient receptor potential vanilloid type 1 (TRPV1), known to react to heat stimuli and capsaicin.

Proinflammatory cytokines sensitize TRPV1 channels, thereby increasing nociceptor excitability and contributing to pain hypersensitivity (Pinho-Ribeiro et al., 2017). During disease or injury, proinflammatory cytokines such as IL-1α, IL-1β, IL-6 and TNF accumulate within the nervous system and, in chronic conditions such as multiple sclerosis (MS) or nervous system trauma, are continuously replenished by recruited immune cells (Bretheau et al., 2022; Levesque et al., 2016; Nadeau et al., 2011; Pineau and Lacroix, 2007), thereby maintaining a state of hypersensitivity (Grace et al., 2014). These cytokines increase nociceptor excitability through activation of their respective receptors (Fukuoka et al., 1994; Junger and Sorkin, 2000; Obreja et al., 2002), in part by increasing sodium channel activity and/or expression and sensitizing TRP channels (Cook et al., 2018), whereas evidence for a direct effect on potassium channels remains limited. This, along with other inflammatory mediators liberated by surrounding cells, was shown to drive central sensitization and chronic pain, leading experts to claim that chronic pain is a neuroinflammatory disorder (Ji et al., 2016).

The excessive production of IL-1 is linked to pathological changes observed in various disorders characterized by pain, including SCI and PNI, as well as chronic inflammatory autoimmune diseases such MS, rheumatoid arthritis (RA), and osteoarthritis (OA) (Boilard et al., 2010; Bretheau et al., 2022; Levesque et al., 2016; Mailhot et al., 2020; Nadeau et al., 2011).

Injecting IL-1α or IL-1β in the cerebrospinal fluid (CSF) or directly into the sciatic nerve was found to induce physiological and behavioral indicators of pain (Nadeau et al., 2011; Zelenka et al., 2005). IL-1 cytokines can also modulate the excitability of nociceptors and their receptors, including TRPV1, GABA, AMPA and NMDA receptors, as well as Nav1.7-1.9 and KCa1.1 channels (Basbaum et al., 2009; Kawasaki et al., 2008; Schafers and Sorkin, 2008; Stemkowski et al., 2021). Further, IL-1 cytokines can increase NGF expression, a neurotrophin recognized as a key mediator of pain (Kusik et al., 2026; Safieh-Garabedian et al., 1995). IL-1β can also trigger intricate signaling pathways, resulting in the release of proalgesic substances such as substance P, bradykinin, and ATP (Economides et al., 2003; Inoue et al., 1999; Samad et al., 2001).

Accordingly, studies have demonstrated that pain-related behaviors can be reduced by neutralization of IL-1R1 through administration of the IL-1 receptor antagonist anakinra, function-blocking antibodies, or using knockout mice (Kwilasz et al., 2021; Mailhot et al., 2020; Ren and Torres, 2009). However, only a limited number of studies have investigated the possibility that IL-1 may stimulate nociceptors directly.

Building on our recent work, in which we revealed that IL-1 cytokines can directly impact a subset of nociceptors and stimulate pain in a variety of chronic inflammatory diseases (Mailhot et al., 2020), we aimed here to characterize the IL-1R1-expressing neurons in the DRGs and their downstream responses to IL-1β stimulation. Using spatial and single-cell transcriptomics, we show that IL-1R1^+^ DRG neurons belong to the NP3 sensory neuron class, typically associated with inflammatory pain and itch, and coexpress markers such as somatostatin (SST) and adenosine deaminase (ADA). Moreover, we found that IL-1β activation of IL-1R1^+^ DRG neurons induces the expression of genes more closely associated with pain than with immune responses.

## RESULTS

### Sex differences in expression of neuronal IL-1R1 in the DRGs of mice

We previously reported that IL-1R1 is expressed by a subset of TRPV1^+^ DRG neurons involved in pain transmission (Mailhot et al., 2020). Important differences have been observed between sexes in terms of pain mechanisms, particularly in relation to the neural circuits and neuroimmune mediators implicated (Mogil, 2020), both of which could be influenced by the IL- 1 cytokine system. To map the distribution of IL-1R1^+^ neurons, we quantified the percentage of NeuN^+^ neurons that were immunostained for IL-1R1 across cervical (C), thoracic (T), and lumbar (L) DRGs in adult male and female C57BL/6 mice. Combining both sexes, we observed spinal level-dependent variation in the abundance of IL-1R1^+^ NeuN^+^ neurons, with the highest counts at C3-C5, T2-10, and L2-L6, peaking at L3-L5 (Fig. 1A). We then compared the average percentage of IL-1R1^+^ NeuN^+^ neurons per DRG between sexes under normal conditions and found that females had ∼175% more IL-1R1^+^ NeuN^+^ neurons than males across all spinal levels.

**Figure 1.**
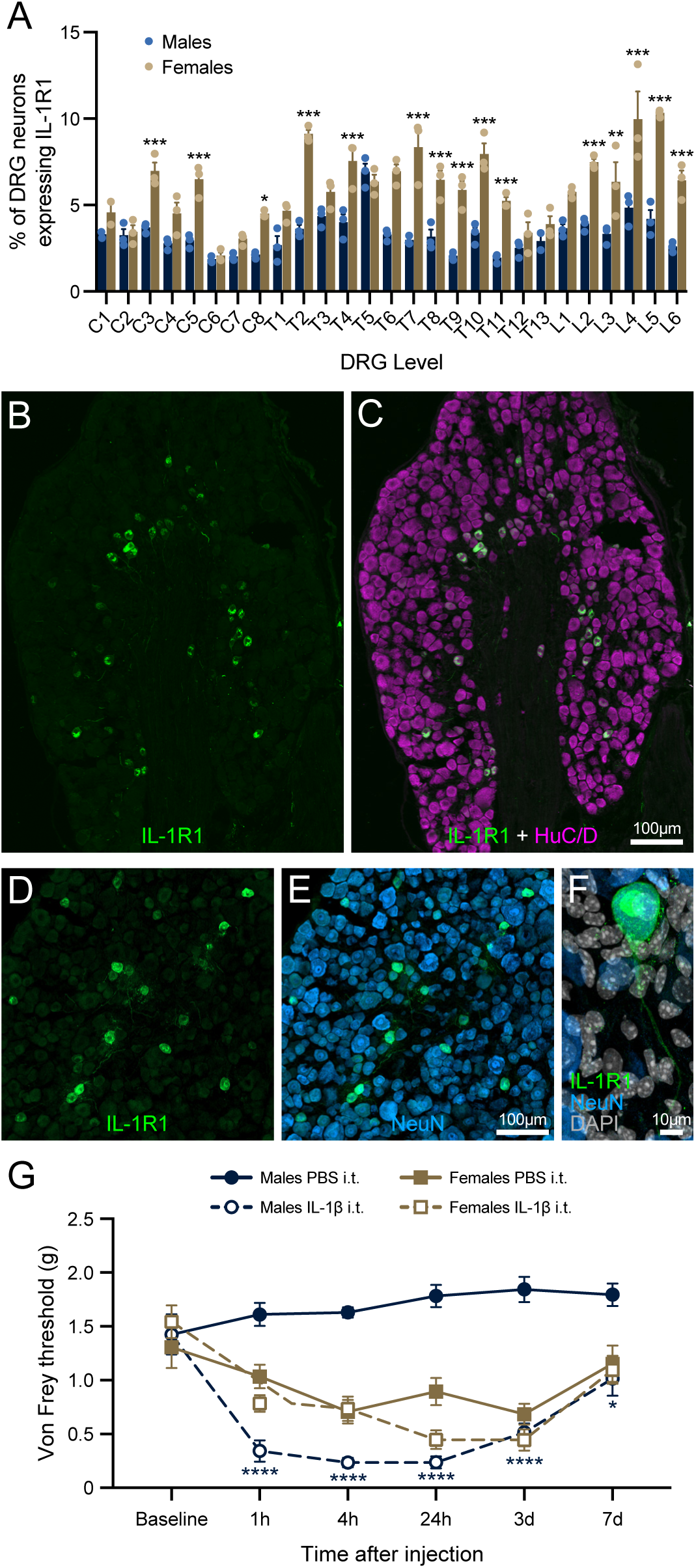
IL-1R1 is expressed by a subset of DRG sensory neurons and CSF IL-1β drives stronger and longer-lasting mechanical pain responses in males than in females. (A) Quantification of the percentage of NeuN^+^ neurons co-expressing IL-1R1 in cervical (C), thoracic (T) and lumbar (L) dorsal root ganglia (n = 6 mice; 3 males and 3 females). (B-C) Representative confocal images showing immunofluorescence for IL-1R1 (green) and the pan-neuronal marker HuC/D (purple) in the L5 DRG. (D-F) Confocal images showing IL-1R1 immunofluorescence localized to the membrane surface of small-diameter NeuN^+^ DRG neurons (D-E), as well as within axons and cytoplasmic regions resembling the endoplasmic reticulum surrounding the DAPI-stained nucleus (F). (G) Mechanical allodynia was assessed using von Frey filaments in male and female mice following intrathecal (i.t.) injection of recombinant mouse IL-1β or PBS (n = 7 mice per sex per treatment). Scale bars: 100 µm in B-C (shown in C), 100 µm in D-E (shown in E), and 10 µm in F.

At the cellular level, IL-1R1 was detected in small-diameter neurons typically located in the inner regions of the ganglion, near nerve fiber bundles (Fig. 1B-C). At the subcellular level, IL-1R1 localized to the cell membrane of these neurons and within the cytosol, primarily associated with perinuclear structures resembling the endoplasmic reticulum, as well as in the axon itself (Fig. 1D-F).

### Male mice exhibit stronger and long-lasting mechanical pain responses than females following IL-1β administration into the CSF

To assess sex differences in IL-1β-induced pain, recombinant mouse IL-1β was injected into the CSF via the intrathecal (i.t.) route in adult male and female C57BL/6 mice, and mechanical allodynia was measured using calibrated von Frey filaments. Although our previous work showed that intraarticular IL-1β injection in the knee joint induces transient allodynia peaking between 4 and 24 hours in C57BL/6 mice (Mailhot et al., 2020), we opted for central administration here to stimulate a broader population of IL-1R1^+^ DRG neurons. As shown in Fig. 1G, baseline mechanical allodynia did not differ significantly between male and female mice.

Injection of PBS into the CSF did not alter mechanical pain thresholds relative to baseline over one week within either sex. However, following central PBS administration, females exhibited significantly greater sensitivity to von Frey filaments than males at all time points (p < 0.0001), indicating a main effect of sex. Intrathecal IL-1β injection elicited stronger and more persistent mechanical allodynia than PBS in males, with effects evident from 1 hour and maintained for up to 7 days post-injection. In contrast, females showed no significant difference in mechanical sensitivity between IL-1β and PBS treatments. Comparison of the two IL-1β groups following i.t. injection revealed a significant main effect of sex (p = 0.0481), with males displaying a lower mechanical threshold than females at 1 and 4 hours, suggesting an earlier onset of inflammatory pain in males. These results indicate that activation of IL-1β/IL-1R1 signaling promotes mechanical allodynia, particularly in male mice.

Taken together, these findings highlight sex-dependent differences in IL-1β-mediated pain signaling in DRG neurons, emphasizing the critical importance of anatomical context and sex as key biological variables in pain research. Notably, males exhibited greater mechanical pain sensitivity than females following IL-1β CSF administration.

### Integrated spatial and single-cell transcriptomics identify the unique molecular signature of IL-1R1⁺ DRG neurons

Having established that IL-1R1 expression in TRPV1^+^ nociceptors contributes to inflammatory pain, we next sought to characterize this subtype of DRG neurons using high-throughput transcriptomic approaches. First, we leveraged NanoString’s GeoMx Digital Spatial Profiling (DSP) platform, which integrates high-resolution immunofluorescence (IF) microscopy with sequencing-based spatial transcriptomics (Merritt et al., 2020), to capture spatially resolved gene expression profiles in DRG tissue (Fig. 2A). Given the high abundance of IL-1R1^+^ neurons in L3-L5 and the stronger IL-1β-induced pain response in male mice, we opted for slide-mounted paraformaldehyde (PFA)-fixed L3-L5 DRG sections from males, while avoiding potential gene expression variability associated with the female estrous cycle. Using the GeoMx Mouse Whole Transcriptome Assay (WTA), we compared gene expression differences between three DRG neuronal populations identified by immunofluorescence based on expression (or lack thereof) of HuC/D (pan-neuronal marker), TRPV1 (nociceptor marker), and IL-1R1: 1) HuC/D^+^ TRPV1^+^

**Figure 2.**
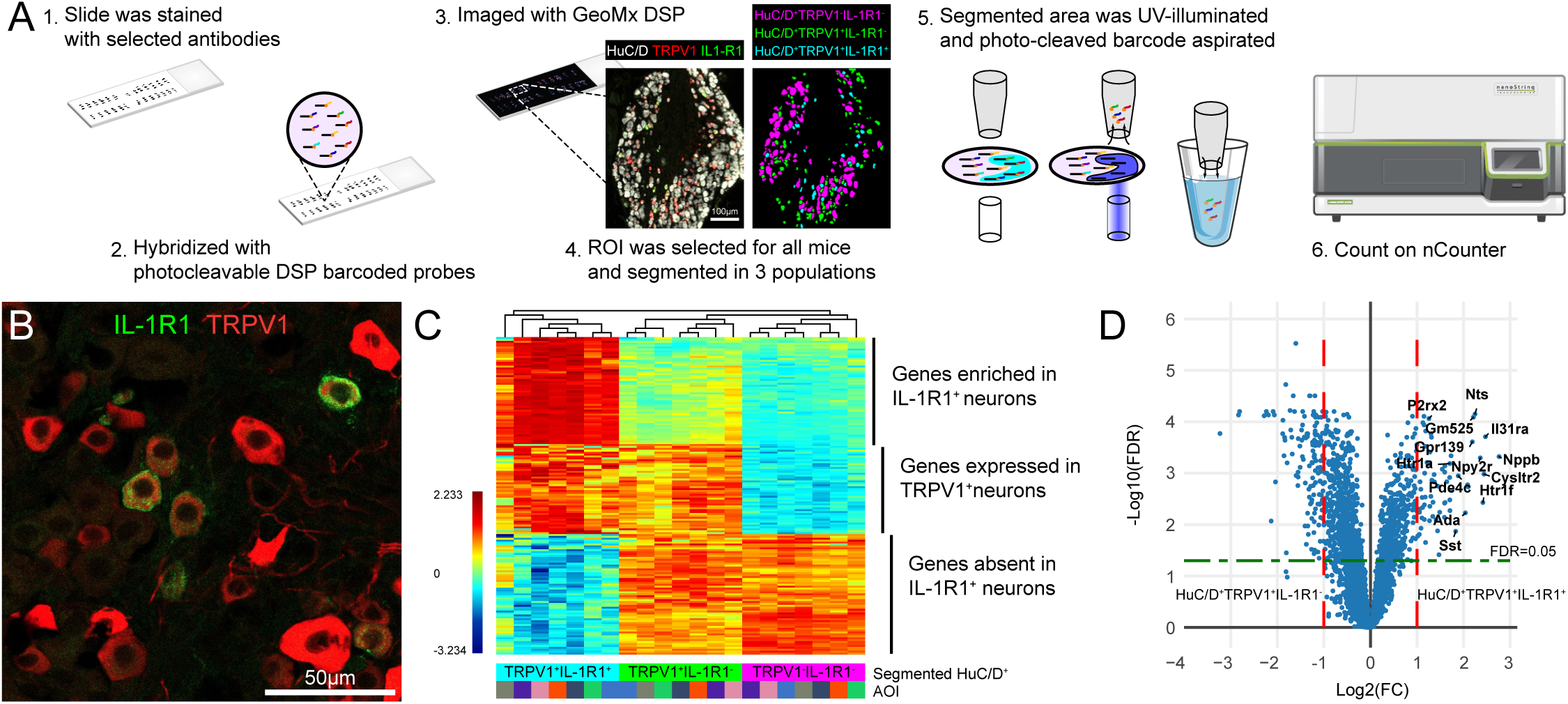
Spatial transcriptomic profiling of IL-1R1-expressing neurons in DRGs reveals their unique gene signature. (**A**) Schematic illustrating the workflow used to profile gene expression in DRG tissue sections via NanoString’s GeoMx Digital Spatial Profiler (DSP). (**B**) Confocal image showing immunofluorescence for IL-1R1 (green) and TRPV1 (red) in an L5 DRG section from an adult naïve C57BL/6 mouse. (**C**) Heatmap of cluster analysis from 21 areas of illumination (AOIs), each corresponding to one of three possible DRG neuron populations in adult naïve C57BL/6 mice: HuC/D^+^ TRPV1^+^ IL-1R1^+^ versus HuC/D^+^ TRPV1^+^ IL-1R1^-^ versus HuC/D^+^ TRPV1^-^ IL-1R1^-^ neurons (n = 7 mice). (**D**) Volcano plot showing differential gene expression between HuC/D^+^ TRPV1^+^ IL-1R1^+^ and HuC/D^+^ TRPV1^+^ IL-1R1^-^ neurons. The green line indicates a false discovery rate (FDR) of 5%, and red lines denote log2 fold change (FC) thresholds. Statistical significance was determined using a t-test with Benjamini-Hochberg correction. Scale bar: 50 µm in **B**. Abbreviation: ROI, region of interest.

IL-1R1^+^ neurons, 2) HuC/D^+^ TRPV1^+^ IL-1R1^-^ neurons, and 3) HuC/D^+^ TRPV1^-^ IL-1R1^-^ neurons (Fig. 2A-C). A spatial transcriptomics approach was chosen given the scarcity of IL-1R1^+^ neurons in DRGs (∼5-10% depending on sex; Fig. 1A), and seeing as we did not know whether *Il1r1* mRNA expression would equate to protein expression. Neuronal populations were selected using contour profiling based on marker expression. In total, the GeoMx Mouse WTA detected between 10,000 and 15,000 genes in at least one of the three DRG neuronal populations. The analysis of spatial transcriptomics data led to the identification of genes that are highly enriched in HuC/D^+^ TRPV1^+^ IL-1R1^+^ DRG neurons compared to IL-1R1-negative nociceptors, such as *Ada, Cysltr2, Gm525, Gpr139, Htr1a, Htr1f, Il31ra*, *Nppb, Npy2r, Nts, P2rx2, Pde4c,* and *Sst* (Fig. 2D). These genes may be involved in the nociceptive effects of IL-1 cytokines or serve as new markers for IL-1R1^+^ nociceptors.

To validate spatial transcriptomic data and further compare gene expression profiles of IL-1R1^+^ neurons with other sensory neuron types, we developed a protocol for efficient isolation of viable neurons from adult mouse DRGs, enabling single-cell RNA sequencing (scRNA-Seq) (Fig. 3A). We used 10x Genomics’ Chromium Single Cell Gene Expression Solution and performed unbiased clustering using uniform manifold approximation and projection (UMAP) analysis (Becht et al., 2018). To characterize IL-1R1^+^ neurons and their response to IL-1β stimulation, we performed scRNA-seq on neurons enriched from pooled C3-C5 and L3-L5 DRGs of male C57BL/6 mice, either naïve or intracisternally (i.c.m.) injected with IL-1β. Mice receiving IL-1β were killed at 1, 4 or 24 hours post-injection to capture temporal changes in gene expression. A total of 14,933 single-cell transcriptomes were analyzed, including 935 cells from naïve mice, 1,753 from the 1-hour IL-1β group, 7,992 cells from the 4-hour IL-1β group, and 4,253 cells from the 24-hour IL-1β group. Approximately 50% of the recovered cells were classified as neurons based on expression of a set of pan-neuronal genes (e.g., *Tubb3*, *Snap25*).

**Figure 3.**
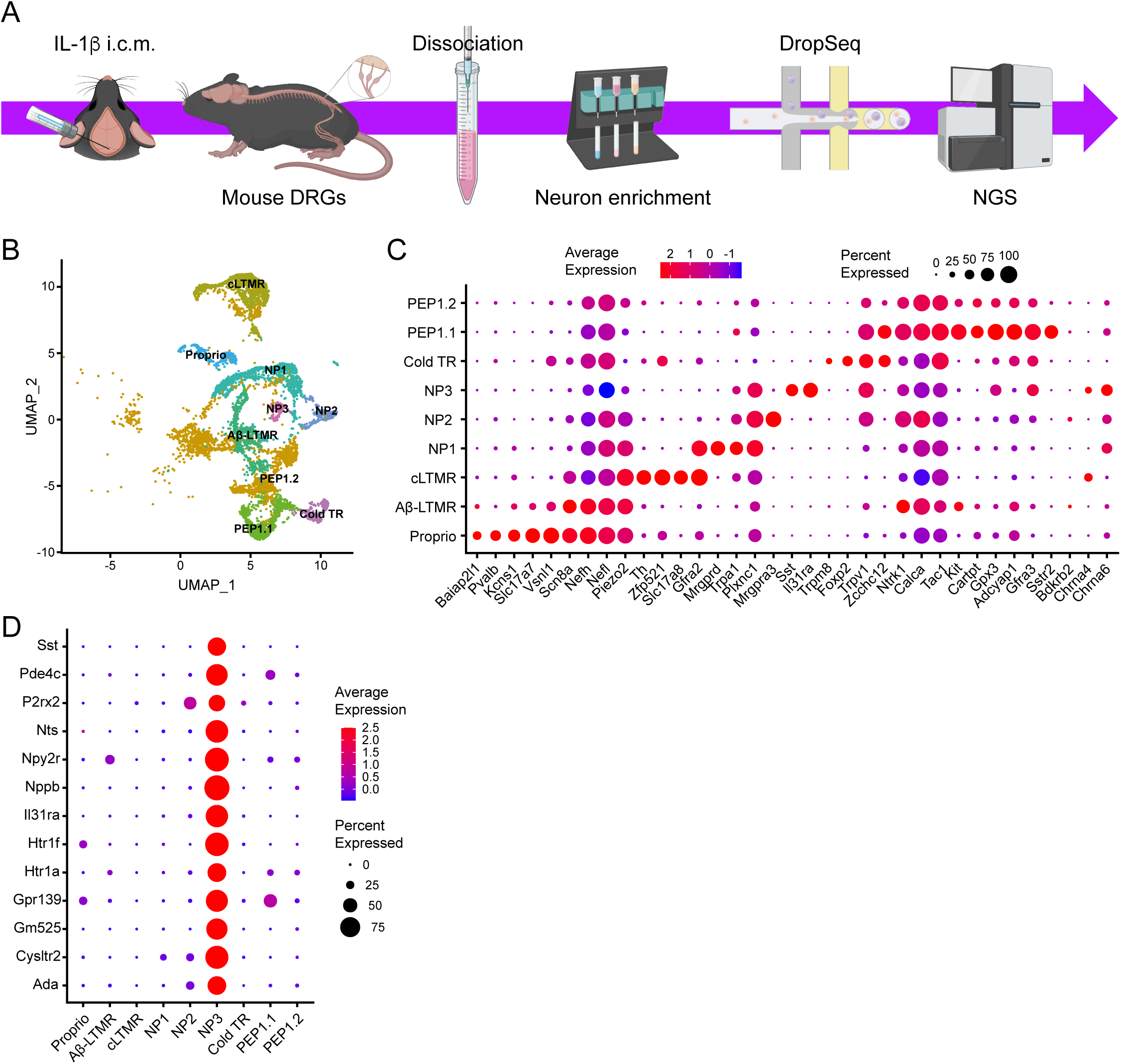
Integration of spatial transcriptomics with single-cell RNA-sequencing of mouse DRGs shows that IL-1R1^+^ neurons belong to the NP3 population which uniquely expresses specific gene markers. (**A**) Schematic of the workflow from adult mouse DRG dissociation to neuronal isolation and enrichment, followed by single-cell RNA sequencing (scRNA-seq) using 10x Genomics DropSeq next-generation sequencing (NGS) technology. (**B**) UMAP-based scatterplot showing nine clusters of DRG neuron types derived from 14,933 freshly isolated cells. (**C-D**) Dot plot analysis showing genes that distinguish DRG sensory neuron types (**C**) and genes specifically marking IL-1R1^+^ nonpeptidergic subtype 3 (NP3) neurons (**D**). The genes shown in panel **D** were selected from spatial transcriptomic data generated using the NanoString platform.

Non-neuronal cells were excluded from analysis based on the expression of established markers for endothelial cells (*Pecam1*, *Cldn5*, *Sox17*), pericytes (*Acta2*, *Rgs5*) fibroblasts (*Ly6a*, *Ly6c1*, *Fabp7*), Schwann cells/satellite cells (*Ncmap*, *Cldn19*), and macrophages (*Fcer1g*, *Cd74*, *Itgam*).

Analysis of scRNA-seq data from recovered DRG neurons across all groups revealed at least nine distinct neuronal clusters (Fig. 3B-C). Each cluster exhibited a comparable number of unique molecular identifiers (UMIs) and an average of ∼2,000 detected genes per cell. Based on UMI expression patterns and established functional profiles from previous scRNA-seq studies (Kupari et al., 2021; Usoskin et al., 2015), these clusters were classified into known DRG sensory neuron subtypes. Mapping marker gene expression onto the UMAP representation revealed the following nine distinct subtypes: peptidergic type 1 (PEP1.1 and PEP1.2), cold thermoreceptors (Cold TR), nonpeptidergic types 1-3 (NP1-NP3), C low-threshold mechanoreceptors (cLTMR), Aβ low-threshold mechanoreceptors (Aβ-LTMR), and proprioceptors (Proprio). Integration of spatial and single-cell transcriptomic data revealed that IL-1R1^+^ DRG neurons correspond to the NP3 sensory neuron cluster, previously associated with inflammation-induced itch (Usoskin et al., 2015). As shown in Fig. 3D, several genes identified as enriched in IL-1R1^+^, but not in IL-1R1-negative, DRG neurons via spatial transcriptomics were found to be relatively specific markers of the NP3 population when cross-referenced with the scRNA-seq dataset. These include *Ada, Cysltr2, Gm525, Gpr139, Htr1a, Htr1f, Il31ra*, *Nppb, Npy2r, Nts, P2rx2, Pde4c,* and *Sst*, all of which showed limited expression in neuronal subtypes other than NP3.

### SST and ADA are relatively specific markers of IL-1R1^+^ DRG neurons

To validate transcriptomic findings and further characterize IL-1R1⁺ neurons, we next examined protein-level expression of candidate genes using IF in mouse DRG tissue. Quantification revealed that approximately 75% of IL-1R1⁺ DRG neurons colocalized with somatostatin (SST), and 90% with adenosine deaminase (ADA), in mouse L5 DRGs (Fig. 4). This contrasts with the 100% colocalization previously observed with TRPV1 in our earlier work (Mailhot et al., 2020). However, consistent with scRNA-seq data shown in Fig. 3C-D, SST and ADA appear to be more specific markers of the IL-1R1⁺ population than TRPV1: ∼50% of SST⁺ and ∼40% of ADA⁺ DRG neurons coexpressed IL-1R1 (Fig. 4D, H), compared to only ∼20% of TRPV1⁺ neurons (Mailhot et al., 2020). These findings validate SST and ADA as more selective comarkers of IL- 1R1⁺ neurons than TRPV1.

**Figure 4.**
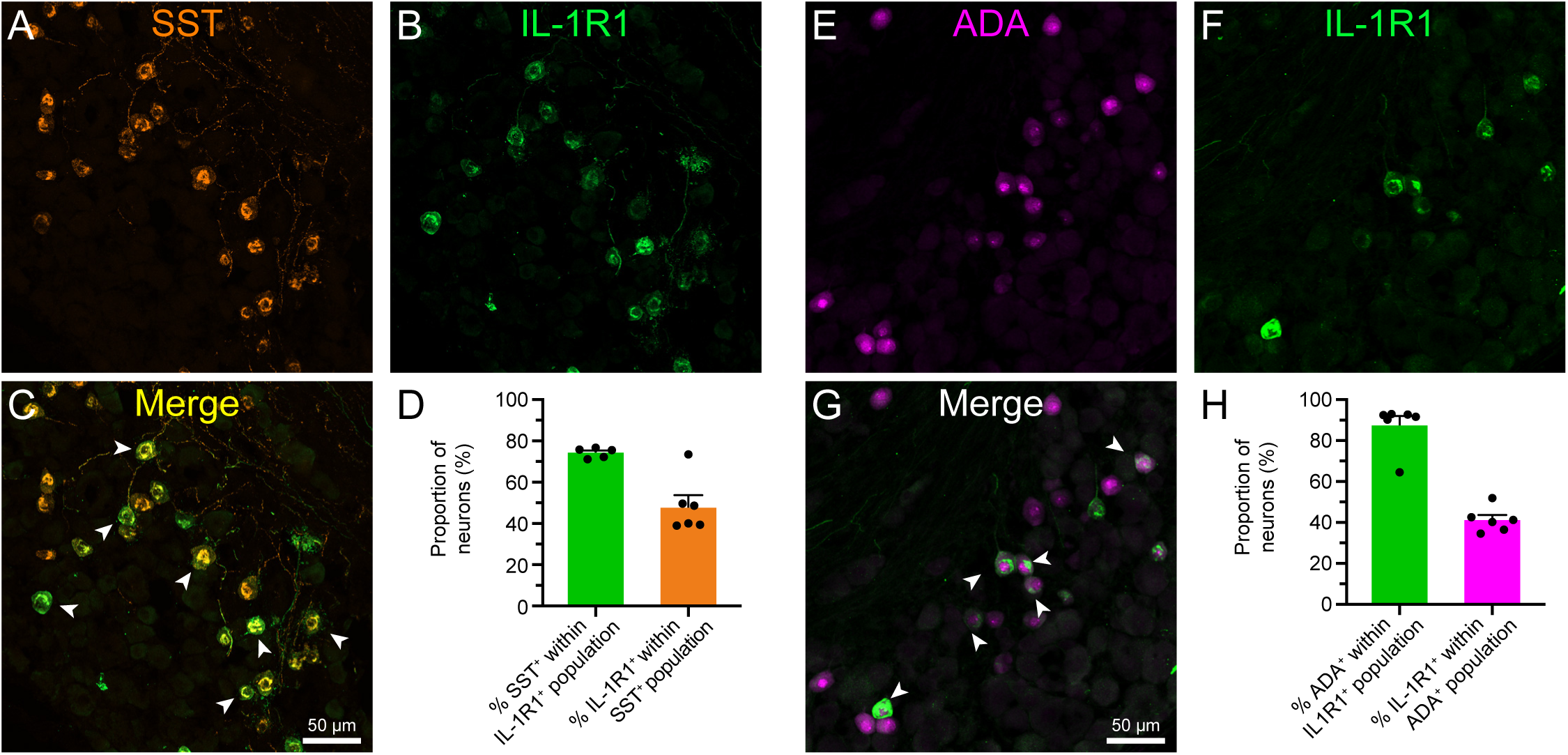
Immunofluorescence validation of somatostatin and adenosine deaminase as specific markers for IL-1R1^+^ DRG neurons in mouse. (**A-C**) Representative confocal images showing somatostatin (SST, orange) and IL-1R1 (green) immunostaining in the L5 DRG of a naïve C57BL/6 mouse. White arrowheads indicate colocalized SST^+^ IL-1R1^+^ neurons in the merged image shown in **C**. (**D**) Quantification of the proportion of SST^+^ neurons within the IL-1R1^+^ neuronal population and of IL-1R1^+^ neurons within the SST^+^ neuronal population (n = 6 mice). (**E-G**) Representative confocal images showing immunostaining for adenosine deaminase (ADA, magenta) and IL-1R1 (green), along with the merged image. Colocalized cells are indicated by white arrowheads. (**H**) Quantification of the proportion of ADA^+^ neurons within the entire IL-1R1^+^ neuronal population and vice-versa (n = 6 mice). Scale bars: 50 µm in **A-C** and **E-G** (shown in **C** and **G**).

### IL-1β injection into the CSF induces a non-classical, non-inflammatory transcriptomic response in NP3 DRG neurons

Having shown that IL-1β administration into the CSF induces mechanical hypersensitivity and that IL-1R1 is selectively expressed by NP3 DRG neurons, we sought to interrogate their gene expression profiles under naïve conditions and following IL-1β stimulation. Compared to naïve controls, NP3 neurons from mice injected i.c.m. with IL-1β exhibited a distinct transcriptional response that diverged from the classical inflammatory pathways observed in endothelial cells, another cell population known to express IL-1R1. Pathway and process enrichment analysis of differentially expressed genes (DEGs) using Metascape revealed that DRG endothelial cells responded to i.c.m. IL-1β injection with significant enrichment of several immune-related genes and pathways (Fig. 5A). These included *TNF signaling pathway* (e.g., *Rela*, *Map3k8*, *Nfkbia*), *Cytokine signaling in immune system* (*Birc3*, *Csf1*, *Osmr*), *Response to IL-1* (*Cxcl2*, *Gsdmd*, *Irak2*), *NF-kappa B signaling pathway* (*Nfkb1*, *Nfkb2*, *Nfkbia*), *Regulation of inflammatory response* (*Cebpb*, *Il17ra*, *Nod2*), *Positive regulation of cell adhesion/migration* (*Icam1*, *Vcam1*, *Sele*), and *Leukocyte activation* (*Cd40*, *Cxcl12*, *Icam1*). In contrast, NP3 neurons showed enrichment for genes associated with ontology terms such as *Sensory organ development* (e.g., *Bdnf*, *Fos*, *Sparc*), *Response to heat* (*Cryab*, *Hspa1a*), and *Regulation of neuronal synaptic plasticity* (*Bdnf*, *Dlg4*, *Vgf*) following IL-1β injection into the CSF (Fig. 5B). We next analyzed DEGs at various time points after IL-1β injection and found that the transcriptional response peaked at 4 hours (Fig. 5C). A comparison between the 1-hour and 4-hour time points revealed a marked upregulation of several genes that are potent inducers of pain responses at 4 hours, including *Alkal2*, *Bdnf*, *Dbi*, *Lcn2*, *Sparc*, and *Timp3* (Fig. 5D and Table 1). The heatmap also highlighted DEGs that were downregulated at 4 hours compared to 1 hour post-injection, including *Fndc5*, *Gper1*, *Nipsnap1*, *Plcb3*, and *Sst,* all of which have previously been associated with pain inhibition. Temporal expression profiling of these pain-associated genes showed rapid yet transient changes in gene expression following IL-1β injection (Fig. 5E). Overall, our scRNA-seq analysis of IL-1β-induced gene expression in mouse DRGs suggests potential direct or indirect interactions between the IL-1β-IL-1R1 pathway and pain-related genes. This unexpected functional diversity underscores the neuromodulatory role of IL-1β in sensory neurons and supports the hypothesis that NP3 neurons, or a subset thereof, may act as an integrative hub for both nociceptive and immune signaling, ultimately shaping pain processing through mechanisms distinct from classical inflammatory pathways.

**Figure 5.**
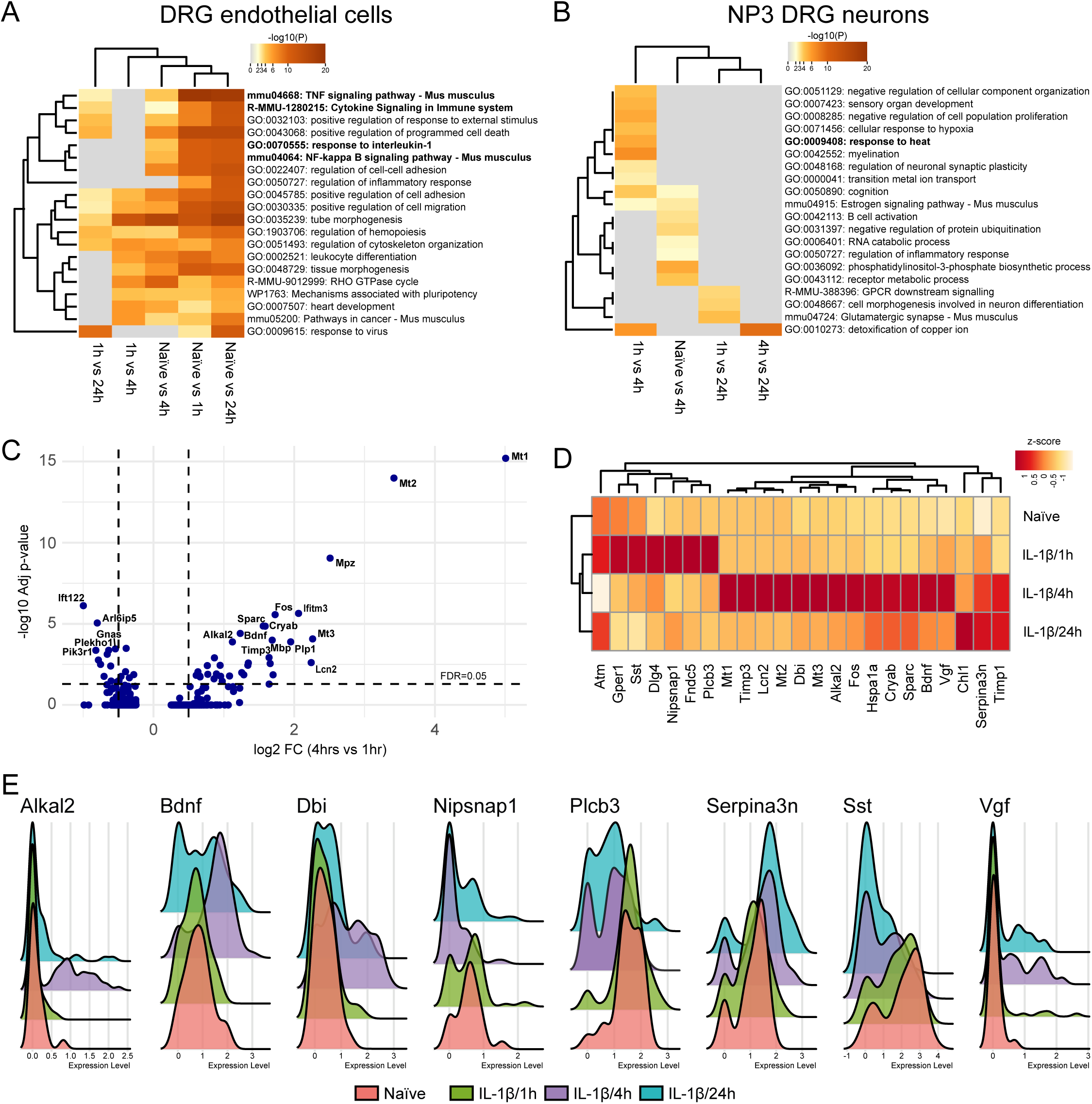
IL-1β injection into the CSF induces a pain-like transcriptional profile in NP3 DRG neurons while promoting an inflammatory signature in endothelial cells. (**A-B**) Heatmaps showing significantly enriched ontology clusters from Metascape analysis of key biological processes associated with DEGs in endothelial cells (**A**) and NP3 neurons (**B**) of DRGs after i.c.m. IL-1β administration compared to naïve mice. (**C**) Volcano plot showing gene expression changes between NP3 neurons 1 hour versus 4 hours post-IL-1β stimulation. The horizontal black line indicates a false discovery rate (FDR) of 5%, whereas the vertical black lines indicate the thresholds of log2 fold change (FC). We assessed significance in differential expression using a t-test with Benjamini-Hochberg correction. (**D**) Heatmap of significantly upregulated and downregulated genes in NP3 DRG neurons under naïve conditions and at 1, 4, and 24 hours (h) following IL-1β injection into the CSF. (**E**) Ridge plots showing the expression of pain-associated DEGs detected in NP3 neurons, such as *Alkal2*, *Bdnf*, *Dbi*, and *Plcb3*, in IL-1β-injected mice at various times points post-injection compared to naïve mice.

**Table 1.**
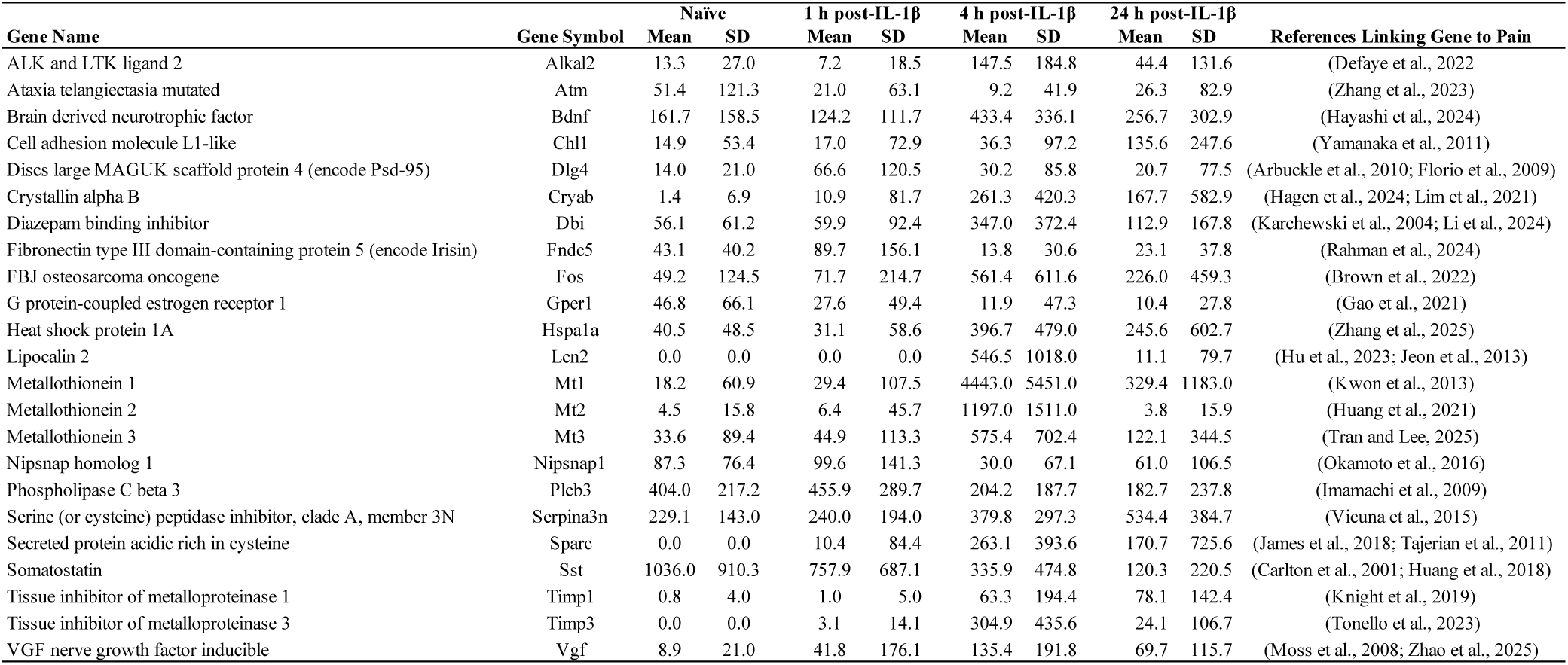
Differentially expressed genes in NP3 DRG neurons following IL-1β administration into the CSF and their roles in pain pathways.

### *Ex vivo* two-photon calcium imaging and behavioral analysis show that IL-1R1-responsive DRG neurons respond to mechanical stimuli and capsaicin, but not to pruritogens

To validate the omics data functionally, we next examined whether IL-1β-responsive neurons in the lumbar DRGs respond to natural nociceptive cutaneous stimuli and pharmacological agents using *ex vivo* two-photon calcium (2P Ca^2+^) imaging. For these recordings, we used an *ex vivo* somatosensory preparation in which the thoracolumbar spinal cord, lumbar DRGs, saphenous and lateral femoral cutaneous nerves, and skin (from the dorsal hind paw to the proximal hip) were dissected in continuity (Fig. 6A). This setup enabled controlled stimulation of the skin with mechanical, thermal (heat or cold), and chemical inputs. We used Snap25-2A-GCaMP6s-D knock-in mice (hereafter Snap25-GCaMP6s), which express the slow-variant calcium indicator GCamp6s in neurons throughout the central nervous system (CNS) and in DRG neurons. Upon calcium binding during neuronal activation, GCaMP6s undergoes a conformational change that increases its fluorescence intensity (Fig. 6B). As a first step in functionally characterizing IL-1β-responsive DRG neurons, we applied recombinant mouse IL-1β (50 nM) to the bath. A responder was defined as a cell showing a stimulus-evoked Ca^2+^ transient with a peak amplitude >50% ΔF/F and >6 standard deviations above baseline. In total, 37 neurons met this criterion and were classified as IL-1β responders out of 2,304 recorded GCaMP6s neurons across three preparations (Fig. 6C). We then assessed functional properties by applying mechanical and thermal stimuli to the skin (Fig. 6D-E). Dynamic mechanical stimulation was delivered using a wide brush, while static stimulation used von Frey filaments (2.0 g for high threshold, 0.16 g for low threshold).

**Figure 6.**
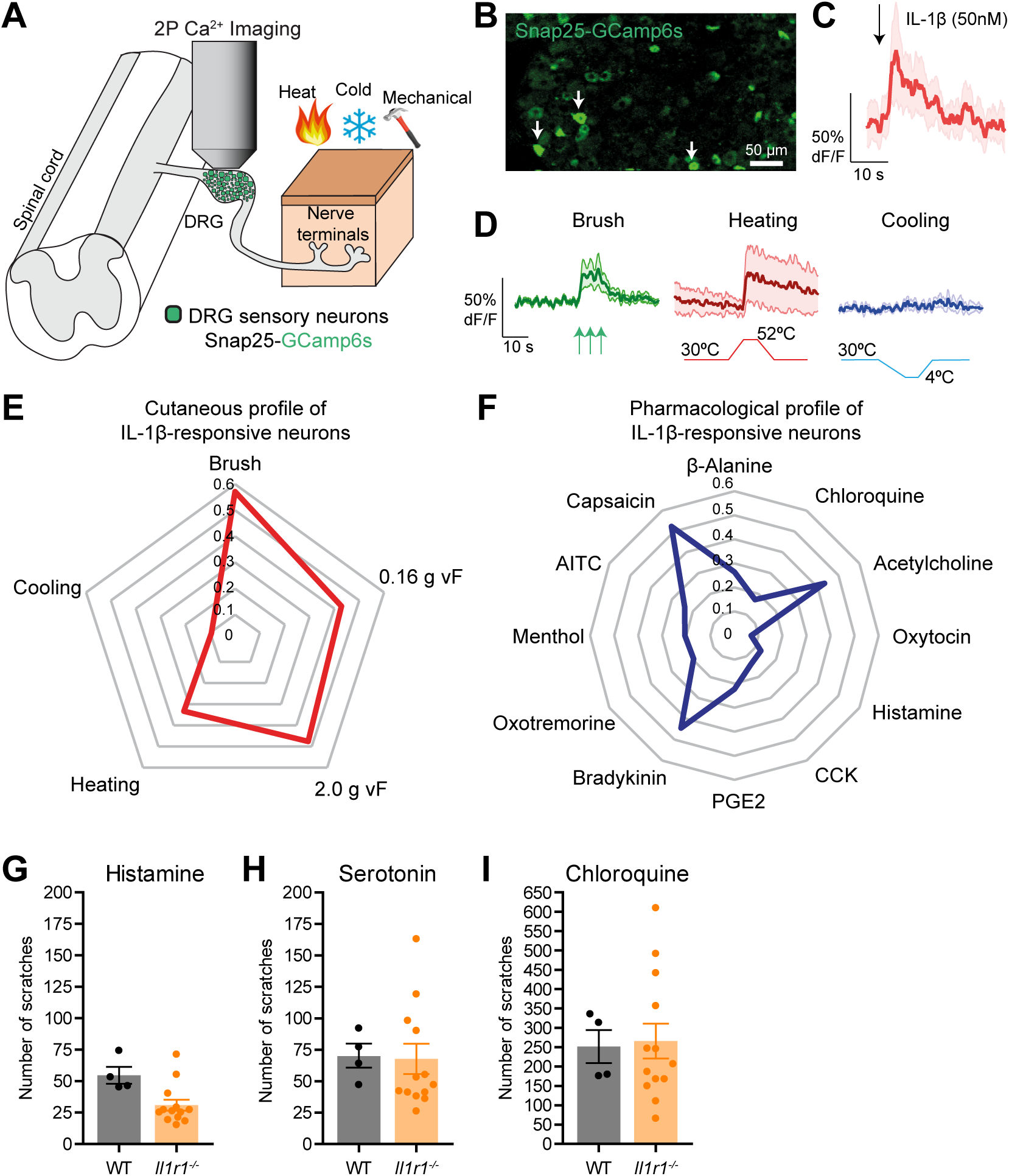
Multiphoton calcium imaging combined with behavioral analysis shows that IL-1β-responsive DRG neurons are activated by cutaneous nociceptive but not pruritogenic stimuli. (**A**) Diagram illustrating the ex vivo preparation and two-photon calcium (2P Ca^2+^) imaging setup for recording DRG sensory neuron responses in Snap25-2A-GCaMP6S-D (Snap25-GCaMP6s) mice to pharmacological and cutaneous stimulation. (**B**) Representative maximum projection of a single imaging plane showing DRG neurons that responded to IL-1β (indicated by arrows) applied through the bath. (**C**) Mean response amplitude of IL-1β-responsive DRG neurons (n = 37 cells, 3 mice). (**D**) Average responses to brushing (green), heating (red), and cooling (blue) the skin. (**E**) Radar chart showing the proportion of IL-1β-responsive neurons also responding to cutaneous stimuli, including punctate mechanical stimulation with von Frey filaments (vF; 2.0 g or 0.16 g). (**F**) Radar chart showing normalized response amplitudes of IL-1β-responsive neurons following application of pharmacological agents to label sensory DRG neuron subtypes. (**G-I**) Number of scratching bouts induced by histamine (**G**), serotonin (**H**), and chloroquine (**I**) after topical application of each pruritogen to the skin of IL-1R1 knockout (*Il1r1*^-/-^) and wild-type (WT) mice (n = 4-13 mice per group). Scale bar: 50 µm in **B**.

Thermal stimuli consisted of heat ramps (30 °C to 52 °C at 4 °C/s) or cold ramps (30°C to 4 °C at the same rate). Among IL-1β-responsive GCaMP6s neurons, ∼60% responded to brush, ∼50% to high-threshold von Frey, ∼45% to low-threshold von Frey, ∼35% to heat, and ∼10% to cold stimulation (Fig. 6E). Thus, many IL-1β-responsive DRG neurons exhibit polymodal sensitivity, responding to multiple sensory modalities and a broad range of mechanical stimuli.

This study, together with our previous work, has established that IL-1R1^+^ DRG neurons express TRPV1, the natural receptor for capsaicin (Caterina et al., 1997), and belong to the NP3 subset, which is associated with inflammatory pain and itch responses. We therefore assessed whether IL-1β-responsive neurons would directly respond to capsaicin and classic pruritogens such as chloroquine and histamine. Among IL-1β-responsive GCaMP6s neurons, approximately 55% responded to bath application of capsaicin, whereas responses to chloroquine (∼20%) and histamine (∼10%) were modest (Fig. 6F). Notably, nearly half (∼45%) of IL-1β-responsive DRG neurons also responded to bradykinin and acetylcholine, suggesting the presence of an NP3 neuron subtype that is largely unresponsive to pruritogens. Because IL-1β-responsive DRG neurons showed minimal responses to bath-applied pruritogens and because itch drives scratching, we next examined scratching behavior in IL-1R1-knockout (*Il1r1*^-/-^) mice. Consistent with 2P Ca^2+^ imaging results, topical application of chloroquine, histamine, and serotonin still evoked robust scratching in *Il1r1*^-/-^ mice, indicating a lack of functional interaction between the IL-1β-IL-1R1 pathway and classical pruritogen receptors (Fig. 6G-I). Collectively, these findings indicate that while a majority of IL-1β-responsive neurons express functional TRPV1, they do not exhibit robust direct responses to itch-causing agents, suggesting a predominant role in inflammatory pain rather than pruritus.

## DISCUSSION

The link between inflammation and pain in disease and injury has long been a subject of intense investigation, yet remains incompletely understood. In this study, we characterized IL-1R1- expressing neurons in the DRGs and investigated their role in IL-1β-mediated inflammatory pain. By integrating spatial transcriptomics with scRNA-Seq, we identified a distinct subpopulation of IL-1R1^+^ sensory neurons in the DRGs, primarily belonging to the NP3 class, which is traditionally associated with inflammatory itch. However, activation of these neurons by IL-1β induced a non-classical transcriptional profile, with gene expression patterns primarily linked to neuronal sensitization and chronic pain, a response that differed from the classical inflammatory profile induced by IL-1β in IL-1R1-expressing CNS endothelial cells. Quantitative analyses revealed sex differences in IL-1R1 expression, with females exhibiting a higher number of IL-1R1^+^ neurons across DRG levels. However, behavioral testing revealed that males developed significantly greater mechanical hypersensitivity than females following IL-1β administration into the CSF.

Our IF data indicate that IL-1R1^+^ DRG neurons are distributed along the rostrocaudal spinal cord axis, with a higher prevalence at the C3-C5 and L2-L6 levels. This pattern suggests that IL-1R1⁺ sensory neurons may be enriched in DRGs innervating regions more exposed to mechanical stress and chronic inflammation, such as the lower back and limbs. Arima and colleagues previously reported that pathogenic CD4^+^ T cells preferentially infiltrate DRGs at these levels in experimental autoimmune encephalomyelitis in mice, a model of neuroinflammation and CNS autoimmunity that recapitulates key features of MS (Arima et al., 2012). Mice with EAE also develop early mechanical allodynia (Olechowski et al., 2009), further highlighting the yet-to-be-defined interactions between inflammation and pain.

We observed that IL-1R1⁺ DRG neurons are significantly more abundant in females than in males. However, despite this higher receptor expression in females, as well as the higher baseline mechanical threshold in males, intrathecal injection of IL-1β induced a more robust and prolonged mechanical hypersensitivity in males than females. This apparent paradox is consistent with previous reports of sex-specific immune and neuronal responses to pain (for review, (Mogil et al., 2024)). Of relevance is the finding that Toll-like receptor 4 (TLR4), a member of the IL-1R/TLR family, mediates hypersensitivity in male but not female mice in both inflammatory (intrathecal LPS) and neuropathic (spared nerve injury) pain models (Sorge et al., 2011). Interestingly, these sex differences were specific to the spinal cord and DRGs, as intracerebroventricular or intraplantar LPS administration produced equivalent allodynia in both sexes. Given that female DRGs express higher IL-1R1 levels than their male counterparts, the sexually dimorphic responses observed may reflect fundamental differences in IL-1β/IL-1R1 signaling cascades or neuroimmune interactions at distinct anatomical levels.

Fan *et al*. elegantly demonstrated that pannexin-1 channels, which interact with components of the inflammasome (such as P2X receptors and caspase-1) and contribute to the regulation of IL-1β production (Silverman et al., 2009), underlie neuropathic pain through distinct cell types and mechanisms in males versus females (Fan et al., 2025). In sciatic nerve-injured male rats and mice, activation of pannexin-1 channels on microglia drives allodynia through the release of vascular endothelial growth factor-A (VEGF-A), whereas in females, pannexin-1 channels on spinal cord-infiltrating CD8^+^ T cells promote mechanical via leptin release (Fan et al., 2025). They further showed that IL–1β neutralization prevented allodynia induced by the adoptive transfer of male-derived pannexin–1-activated microglia, indicating that IL–1β and VEGF-A likely act in a common downstream pain signaling pathway. Another study identified microglial P2X4 receptors, which are also involved in inflammasome activation and IL-1β release (de Rivero Vaccari et al., 2012), as a key point of divergence underlying injury-induced hypersensitivity in male rats compared to females (Mapplebeck et al., 2018). Taken together, these findings provide a clear precedent for why the IL-1β/IL-1R1 axis may differ in its capacity to induce neuropathic pain in males compared to females.

We previously reported that IL-1R1-expressing neurons represent approximately 5-10% of the total neuronal population in the DRGs, and that all of these neurons coexpress TRPV1.

However, the opposite is not true, with only about 20% of TRPV1^+^ neurons expressing IL-1R1^+^ (Mailhot et al., 2020), highlighting the need to identify more specific comarkers. Spatial transcriptomics performed using the GeoMx DSP technology enabled high-resolution classification of DRG neurons based on HuC/D, TRPV1, and IL-1R1 expression. When compared with consensus classifications of mouse DRG neuron types across scRNA-seq datasets and nomenclatures (Kupari et al., 2021), we found that IL-1R1^+^ DRG neurons map to the NP3 class. The assignment of IL-1R1^+^ DRG neurons to a subset of NP3 nociceptors was further confirmed at the protein level by colocalization of IL-1R1 with canonical nociceptor markers (e.g., TRPV1, P2X3) and NP3-selective markers (e.g., SST, ADA) in both mouse and human DRGs, as shown here and in our previous work (Mailhot et al., 2020). Notably, both SST and ADA have previously been shown to suppress mechanical hypersensitivity in models of inflammatory pain (Carlton et al., 2001; Hu et al., 2016). These markers of IL-1R1^+^ DRG neurons are not only promising targets for therapy in IL-1β-driven chronic pain, but also provide tools for more precise genetic targeting of this neuronal population by leveraging these genes as drivers in viral vectors and transgenic mouse models.

Following IL-1β stimulation, IL-1R1^+^ NP3 neurons display a non-canonical transcriptional program that diverges from classical inflammatory pathways, marked by upregulation of genes involved in sensory organ development, response to heat, cellular response to hypoxia, and regulation of neuronal synaptic plasticity. Key IL-1β-induced genes in IL-1R1^+^ NP3 neurons code for: BDNF, a well-established modulator of synaptic plasticity and central sensitization (Hayashi et al., 2024), frequently associated with VGF in pain-related contexts (Zhao et al., 2025); ALKAL2, a ligand for the ALK receptor tyrosine kinase implicated in neuron excitability and pain sensitization (Defaye et al., 2022); TIMP3 and SPARC, which regulate extracellular matrix remodeling and glial-neuronal interactions in pain processing (James et al., 2018; Tajerian et al., 2011; Tonello et al., 2023); and DBI, involved in GABAergic modulation and linked to altered neuronal excitability and peripheral pain gating (Karchewski et al., 2004; Li et al., 2024). The observed upregulation of genes encoding metallothioneins 1-3 (MT1-3) and CRYAB hours after IL-1β injection into the CSF suggests the activation of neuroprotective, immunomodulatory, and antioxidant pathways (Hagen et al., 2024; Huang et al., 2021; Kwon et al., 2013; Lim et al., 2021; Tran and Lee, 2025). Furthermore, LCN2 and SerpinA3N, both induced by IL-1β stimulation, contribute to a neuroimmune axis linking PNI to central sensitization and neuropathic pain (Hu et al., 2023; Jeon et al., 2013; Vicuna et al., 2015). All this supports the idea that NP3 neurons activate a specialized transcriptional program that integrates immune signaling with neuronal remodeling, potentially driving pain chronification through mechanisms distinct from classical inflammatory pathways.

The NP3 cluster also expresses *Osmr* and *Il31ra*, which together form the receptor for IL- 31, a cytokine implicated in itch-related inflammatory conditions such as atopic dermatitis (Sonkoly et al., 2006). Pain and itch mechanisms are closely intertwined and share several molecular mediators, including the neuropeptide SST (Huang et al., 2018). However, our data show that responses to classical pruritogens such as histamine, serotonin, and chloroquine are not altered in *Il1r1*^-/-^ mice, suggesting that IL-1R1 may be involved in other forms of inflammatory pain rather than classical itch pathways.

Comparative transcriptomic analyses of mouse, primate, and human DRG neurons have revealed a high degree of conservation in neuronal identities across species (Kupari et al., 2021), suggesting that translating findings from rodents to humans is a realistic goal. Major sensory neuron subclasses identified in mouse DRGs, including peptidergic, non-peptidergic and proprioceptive populations, are transcriptionally conserved in primates and humans, with shared marker genes and core molecular programs. While species-specific differences exist, particularly in gene expression levels and the representation of certain subpopulations, the overall organization of DRG neuron types appears largely preserved. Extending these findings, recent work from Ted Price and colleagues further supports the translational relevance of rodent models by showing strong cross-species conservation of pain-relevant transcriptional signatures and signaling pathways in human DRG neurons (Bhuiyan et al., 2025). Together, these studies suggest that fundamental mechanisms governing sensory neuron identity and nociceptive signaling are shared across mammals, supporting the notion that insights gained from rodent DRG transcriptomics can be meaningfully translated to human pain biology.

*Ex vivo* two-photon calcium imaging shows that IL-1R1^+^ NP3 neurons define a distinct inflammatory pain-responsive state rather than a broadly sensitized nociceptor population. These neurons respond to IL-1β but not to classical pruritogens, supporting the idea that inflammatory pain and itch are mediated by distinct neuronal populations. Consistent with their expression of TRPV1 and NP3 identity, they are activated by dynamic and static mechanical stimuli, noxious heat, and capsaicin. However, IL-1R1^+^ NP3 neurons were unresponsive to PGE_2_, indicating that cytokine-responsive nociceptors can operate independently of cyclooxygenase-derived mediators. Future work should determine whether NP3 neurons express full-length IL-1R1 or truncated isoforms (τIL-1R1) described in CNS inflammation (Bretheau et al., 2022; Mailhot et al., 2020), which could explain the non-classical, non-inflammatory transcriptional program observed following central IL-1β exposure.

In summary, we identified a distinct population of IL-1R1-expressing nociceptors within the DRGs that are molecularly and functionally distinct from classical peptidergic and mechanosensitive subtypes. These IL-1R1^+^ neurons belong to the NP3 subclass and, upon IL-1β stimulation, exhibit a unique transcriptional signature enriched for genes involved in neuroplasticity, neuronal remodeling, and pain processing, rather than the canonical inflammatory pathways typically activated in IL-1β-stimulated endothelial cells. Using multimodal transcriptomics and calcium imaging, we demonstrated that IL-1β directly activates this neuronal population in a receptor-dependent manner. Importantly, this activation induces mechanical hypersensitivity *in vivo*, with a more pronounced effect in male mice. Furthermore, we uncover potentially novel neuron-intrinsic mechanisms underlying IL-1β-driven pain and provide a cellular basis for the observed sexual dimorphism in inflammatory pain responses. Targeting IL-1R1^+^ NP3 neurons may represent a promising, sex-informed therapeutic strategy for chronic pain conditions associated with inflammation.

## MATERIALS AND METHODS

### Mice

Male and/or female C57BL/6 mice were purchased from Charles River Laboratories or The Jackson Laboratory (JAX) at 8-12 weeks of age. Snap25-GCaMP6s mice (stock #025111) were purchased from JAX and bred and maintained as homozygotes on a C57BL/6 background at the University of Pittsburgh. IL-1R1-KO mice were obtained from Dr. Ning Quan (Florida Atlantic University, Jupiter, FL) (Liu et al., 2015), and subsequently bred in-house at the Animal Research Facility of the Centre de recherche du CHU de Québec–Université Laval. All mice were housed in individually ventilated cages on racks connected to a central HEPA filtered air supply (30-70 air changes per hour), with social housing limited to a maximum of 5 adult mice per cage. Animals were maintained on a standard 12-h light/dark cycle and had free access to food and water. Room temperature was maintained at 23 ± 2 °C with a relative humidity of 50 ± 5%. All procedures were approved either by the *Comité de protection des animaux de l’Université Laval* (CPAUL, protocol #CHU-24-1634) or the University of Pittsburgh Institutional Animal Care and Use Committee (IACUC, protocol #24105512) and complied with the ethical guidelines of the Canadian Council on Animal Care or U.S. NIH Guide for the Care and Use of Laboratory Animals.

### Intra-cerebrospinal fluid injections

Mice were anesthetized with isoflurane and injected either intrathecally (i.t.) or intra-cisterna magna (i.c.m.) with recombinant mouse (rm) IL-1β (100 ng/µl diluted in PBS, 5 µl/mouse; Peprotech, catalog #211-11B) or PBS (5 µl/mouse). Intrathecal injections of either rmIL-1β or PBS were performed at the L5-L6 intravertebral space using 0.5-ml syringes fitted with permanently attached 27-gauge needles (BD Medical, cat. #305620). The i.c.m. treatment consisted of a single injection directly into the CSF beneath the skull, penetrating the dura with a pulled-glass micropipette connected to a 10-µL Hamilton syringe, following our published method (Bretheau et al., 2022).

### Tissue processing

#### Mouse tissues

Mice were overdosed with a mixture of ketamine and xylazine and transcardially perfused with phosphate-buffered saline (PBS) followed by 1% paraformaldehyde (PFA) in PBS (pH 7.4).

After perfusion with the fixative, DRGs were dissected out, post-fixed for 24 hours in 1% PFA at 4°C, and then placed in a PBS/20% sucrose solution until tissue processing. On the day of tissue sectioning, DRGs were embedded in M-1 Embedding Matrix (Thermo Fisher Scientific) and then processed using a cryostat (Leica Microsystems, model CM3050S). DRGs were cut transversely at a thickness of either 8 µm (for spatial transcriptomics) or 12 µm (for immunofluorescence). Tissue sections were collected directly onto Surgipath X-tra® slides (Leica Biosystems), separated into four series of adjacent sections, and then stored at −20°C until use.

#### Immunofluorescence and quantification

Tissue sections were rinsed with PBS, blocked with 10% normal donkey serum in PBS containing 0.25% Triton-X, and incubated overnight at 4°C with the following primary antibodies (dilutions, clone numbers, sources, and catalog numbers indicated in parentheses): rabbit anti-ADA (1:10 dilution; Developmental Studies Hybridoma Bank, CPTC-ADA-1 hybridoma was deposited at the DSHB by Dr. A.P Paulovich), mouse anti-HuC/D (1:80; clone 16A11; Thermo Fisher Scientific, A-21271), goat anti-IL-1R1 (1:100; R&D Systems; AF-771), rabbit anti-NeuN (1:1,000; clone D3S3I; Cell Signaling; 12943), rabbit anti-SST (1:200; Thermo Fisher Scientific, PA5-82678), rabbit anti-TRPV1 (1:250; Alomone Labs; ACC-030), and rabbit anti-TRPV1 (1:2,000; Synaptic Systems GmbH; 444 003). Primary antibodies were visualized following a 2.5 h incubation a room temperature with the appropriate Alexa Fluor®-conjugated secondary antibodies (1:250 dilution; Thermo Fisher Scientific). DAPI (1 μg/ml; Thermo Fisher Scientific) was used for nuclear counterstaining. Sections were imaged on a Zeiss LSM800 confocal microscope system equipped with 405, 488, 561, and 640nm lasers. Confocal images were acquired and mosaics created using the Zen Blue Edition software (v. 2.3, Carl Zeiss).

Confocal stacks were taken at either 10×, 20× or 63× magnification, and processed as maximum intensity projections of confocal z-stacks using the ImageJ software (v. 1.44o, Wayne Rasband, NIH).

Quantification of cells stained by IF was performed using BIOQUANT Life Science software (v. 18.5, Bioquant Image Analysis Corporation). The total number of IL-1R1^+^NeuN^+^ and NeuN^+^ neurons per cross-section was counted at 20× magnification using live imaging.

Similarly, IL-1R1-expressing neurons co-expressing ADA or SST (and vice-versa) were counted live at the same magnification. Only immunolabeled cells with a DAPI-stained nucleus were counted, and results were expressed as the proportion of immunolabeled neurons per mm². All quantifications were done blind with respect to the identity of the animals.

### Mechanical sensitivity assessment

Mechanical allodynia was assessed by stimulating the hindpaw plantar surface with calibrated von Frey filaments (Stoelting) on an elevated wire-mesh grid, measuring pain responses (e.g., paw flinching, rapid withdrawal, or intense licking) during or after filament removal, as described previously (Olechowski et al., 2009). In brief, animals were habituated to the testing environment daily for at least a week before baseline testing. Before experimentation, mice were allowed to acclimatize for at least 1 h in individual Plexiglass chambers with a wire mesh floor (Ugo Basile®). Mechanical sensitivity was assessed using calibrated von Frey filaments (0.04 to 6.0 g) applied to the plantar surface of each hind paw by a blinded experimenter. Paw withdrawal thresholds were determined using the conventional Chaplan up–down method (Chaplan et al., 1994), beginning with the filament 0.6 g. The 50% paw withdrawal threshold was calculated based on the pattern of responses to the sequential von Frey filament applications.

### Acute itch studies

At least 24 hours prior to testing, mice were shaved at the nape of the neck under isoflurane anesthesia. The following pruritogens were dissolved in saline and injected subcutaneously (100 μl) using a 26 1/2-gauge needle into the nape of the neck of *Il1r1*^-/-^ mice and their wild-type (WT) littermates: chloroquine (100 µg, Millipore Sigma, Cat. # C6628), histamine (500 μg, Sigma-Aldrich, Cat. # H7250, and serotonin (100 nmol, Sigma-Aldrich, Cat. #H9523). After acclimatization to the test chamber for 30 min. The resultant bouts of scratching were counted at 5-min intervals over a 30-min observation period.

### Digital spatial transcriptomic profiling

Spatial transcriptomic analyses were conducted with expert support from NanoString’s Technology Access Program team using the GeoMx® DSP platform and the Mouse Whole Transcriptome Atlas (MuWTA, Brucker Spatial Biology). A single slide was prepared by mounting 8-μm-thick L5 DRG tissue sections from naïve mice (n=12) perfused transcardially with 4% PFA. The slide was baked at 60°C for 30 min in a drying oven. RNA targets were exposed and antigens retrieved in 1x Tris-EDTA pH 9.0 at 99°C for 20 min, followed by washes in 1x PBS and incubation with 0.1 μg/mL proteinase K in PBS for 15 min at 37°C. The slide was then incubated overnight at 37°C with RNA probes included in the GeoMx® MuWTA kit (NanoString), conjugated to unique DNA-indexing oligonucleotides (DSP barcodes) and covering >21,000 transcripts, in the provided Buffer R using a humidity chamber. The next day, the slide was washed in stringent SSC washes of 5-15 min each at 37°C, followed by PBS washes and IF using a cocktail of prevalidated antibodies, including mouse anti-HuC/D (Thermo Fisher Scientific), goat anti-IL-1R1 (R&D Systems) and rabbit anti-TRPV1 (Synaptic Systems GmbH), following our above-described IF protocol and Buffer W provided in the GeoMx® kit. The slide was loaded onto the GeoMx® DSP instrument and scanned to obtain images of all DRG sections. Eight regions of interest (ROIs), each measuring 677 µm × 785 µm and nearly covering an entire L5 DRG from different naïve mice (n=8), were selected across the whole slide based on fluorescence intensity and density of IL-1R1^+^TRPV1^+^HuC/D^+^ neurons (i.e., >15 IL- 1R1^+^ neurons). Each ROI was segmented into three areas-of-illumination (AOI, n=24): 1) HuC/D^+^TRPV1^+^IL-1R1^+^ neurons, 2) HuC/D^+^TRPV1^+^IL-1R1^-^ neurons, and 3) HuC/D^+^TRPV1^-^ IL-1R1^-^ neurons. Images of registered DRGs were imported into Fiji (ImageJ) to generate precise masks based on fluorescent profiles using the Threshold, Erode/Dilate, Eraser, and Image Calculator tools, enabling UV illumination of specific neuronal subsets in each AOI. The AOIs were then sequentially exposed to 385 nm light, DSP barcodes were released into solution from the hybridized RNA probes, aspirated into a collection plate well, and dried down for 1 h at 65°C. Each well contained the photocleaved DNA oligo tags of a single AOI. DNA oligo tags were then resuspended in 10 μL nuclease-free water and hybridized to GeoMX® Hyb Codes at 65°C for 18 hours. Samples were finally pooled and processed on NanoString’s nCounter MAX system (Brucker Spatial Biology).

Data analysis was conducted using NanoString’s GeoMx NGS pipeline (v. 2.1). nCounter counts were converted to digital count files and imported into the GeoMx DSP platform to generate an expression count matrix. After ROI quality control per NanoString’s recommendations and principal component analysis (PCA), one outlier ROI was removed, leaving 21 AOI raw counts for full quantile normalization. Differentially expressed genes (DEGs) were tested using the *limma* package, with *p*-values adjusted by the Benjamini-Hochberg method.

### Single-cell RNA sequencing and analysis

#### DRG neuron isolation and enrichment

Mice were anesthetized, killed by decapitation, and their cervical (C3-C5) and lumbar (L3-L5) DRGs dissected out. DRGs were incubated at 37°C for 1 h 45 min in prewarmed 0.2% Collagenase I in DMEM, followed by 15 min in 0.25% Trypsin-EDTA. Tissue was dissociated using a 0.5% BSA-coated 22G needle and filtered through a 70-μm cell stainer. Non-neuronal cells were depleted using the Adult Neuron Isolation Kit (Miltenyi Biotec).

#### Single-cell RNA library preparation and sequencing

Approximately 8,700 cells per sample were loaded into a Chromium Single Cell Chip (10x Genomics). Reverse transcription and library preparation were performed using the Chromium Single Cell 3’ Library and Gel Bead Kit v3 (10x Genomics) per manufacturer’s instructions.

Samples were sequenced to an average depth of 43,000 reads per cell on an Illumina NovaSeq sequencer. Cell Ranger (10x Genomics, v. 3.0.1) was used for demultiplexing, barcode processing, unique molecular identifiers (UMI) filtering, gene counting, and mapping to the *Mus musculus* (mouse) reference transcriptome from the Genome Reference Consortium Mouse Build 38 (GRCm38/mm10).

#### Analysis of scRNA-seq data

Raw sequencing data were converted to expression count matrices using the Cell Ranger software (Single Cell Software Suite, 10x Genomics, v. 7.1) and aligned to the GRCm38/mm10 mouse reference genome. The Seurat method was applied to downstream analysis (Satija et al., 2015). Cells with fewer than 1,000 molecules were excluded, and genes expressed in at least five cells with expression levels >1 were considered. A total of 14,933 cells met the criteria. Variable genes were identified using the “vst” method for PCA. Cluster markers were identified using the FindAllMakers function in Seurat (v. 4.3), applying a two-sided Wilcoxon rank-sum test with Bonferroni correction for multiple comparisons. Significant PCs were used for dimension reduction via UMAP to identify cell clusters with DEGs. Genes with absolute log fold change > 0.5 and FDR *p*-value < 0.05 were selected as DEGs.

#### Comparison of transcriptomic profiles of DRG cells after IL-1β stimulation

To compare global expression profiles of DRG cells under IL-1β stimulation, cell counts from each scRNA-seq sample were aggregated to generate pseudo-bulk samples, which were analyzed using bulk RNA-Seq methods. Genes with low expression (average read count < 10) were excluded, resulting in a dataset of 16,059 genes. Raw counts were normalized using the trimmed mean of M-values (TMM) algorithm in edgeR, and transformed to log2 counts per million (logCPM) using the voom function in *limma*. DEGs were identified using the Imfit function, with *p*-values adjusted for multiple testing using the Benjamini-Hochberg method. Pearson correlation coefficients of logCPM values were calculated using the cor function in R to create a correlation heatmap. Pathway and process enrichment analysis was conducted using Metascape (v. 3.5), an integrated web-based portal for functional enrichment and interactome analysis leveraging over 40 independent knowledgebases (Zhou et al., 2019). Gene Ontology (GO) biological processes, Reactome, KEGG, WikiPathways were selected, and default parameters were applied using the multiple gene list tool on a pre-ranked list of DEGs filtered by fold change (FC ≥ 2.0 or ≤ −2.0) and adjusted *p* value < 0.05.

### Two-photon calcium imaging of DRG sensory neurons

#### Ex vivo skin nerve preparation

Mice were deeply anesthetized with ketamine/xylazine, and the right hind paw, leg, and back shaved with an electric clipper, leaving 2-3 mm of hair. Mice were then transcardially perfused with ice cold sucrose-based artificial cerebrospinal fluid (aCSF) saturated with 95% O_2_/5% CO_2_. The sucrose-based aCSF contained (in mM): 234 sucrose, 2.5 KCl, 0.5 CaCl_2_, 10 MgSO_4_, 1.25 NaH_2_PO_4_, 26 NaHCO_3_, and 11 glucose. A dorsal laminectomy was performed, followed by removal of the spinal column, right ribs, and right leg. Tissue was placed in a dissection dish, and the lateral femoral cutaneous and saphenous nerves were dissected out in continuum from the skin to the DRG. The thoracolumbar spinal cord was removed from bone, roots gently separated, and dura removed. The DRG was left in the spinal foramen and the vertebral column stabilized with Minutien pins (Fine Science Tools, catalog # 26002-20) on a Sylgard-coated chamber (Dow Corning, Sylgard 184). Skin was pinned to a metal platform away from the DRG to allow objective placement and imaging without interfering with skin stimulation. The recording dish was then transferred to the microscope and perfused with normal aCSF (in mM: 117 NaCl, 3.6 KCl, 2.5 CaCl_2_, 1.2 MgCl_2_, 1.2 NaH_2_PO_4_, 25 NaHCO_3_, 11 glucose) over 15 min. The temperature of the recording solution was slowly raised to 28°C, 1L of normal aCSF was washed over the preparation to washout any remaining sucrose, and the remaining 4L recirculated for the remainder of the experiments.

#### Multiphoton calcium imaging

Imaging was performed as previously described with modifications to image the DRG instead of the spinal cord (Sheahan et al., 2024; Warwick et al., 2022). Calcium (Ca^2+^) imaging was performed on a ThorLabs Bergamo II multiphoton microscope using ThorImage (v. 4.3) software and a Nikon 16×, 0.8 NA, 3.0 mm WD water-dipping objective. The microscope was equipped with an 8 kHz Galvo-Resonant Scanner, Piezo Objective Scanner, and two GaAsP detectors. The imaging light was provided by a Spectra-Physics Mai Tai HP-244 laser tuned to 940 nm. Emission light was collected through a 570 nm dichroic mirror and FITC filter. The DRG was imaged across five imaging planes spaced 25 µm apart (100 µm depth). Imaging was performed at 60 Hz for the total volume and ∼8 Hz per plane over a 512 x 256 pixels field. Magnification was set to ∼1.4 (pixel size ∼1 µm). This approach enabled imaging of ∼400-500 DRG neurons per experiment.

A mechanical search stimulus in the form of a wide brush was used to identify the region of skin with the greatest cutaneous input. Once identified, this region was used for all subsequent stimulation, including 2.0 g and 0.16 g von Frey hairs applied over a 15 mm x 15 mm area of skin in a 4 x 4 grid. All natural stimuli were applied to the same 15 x 15 mm area using stimulators designed to fit in the same footprint. A dim red light was focused on the skin to aid the experimenter and minimize sensor light contamination. For manual stimuli, a program was written in Signal (CED) to generate synchronized audio cues: a 3-2-1 countdown followed by a 1 s tone indicating stimulus duration, ensuring consistent timing.

For thermal stimulation, a custom-made water-cooled Peltier was placed on the region of skin and held at 30 °C between trials. The Peltier was driven by a custom-designed controller to the indicated temperatures at specified rates and set point. Surface temperatures were recorded in Signal 7 (CED) via a CED 1401. Heat ramps increased from 30 °C to 52 °C at 4 °C/s, held for 5 s, then returned to baseline. Cold ramps decreased from 30°C to 4 °C at the same rate. All thermal stimuli were applied three times for each testing block separated by at least 1 min.

All imaging, stimulus controllers, feedback sensors (e.g., thermocouples), and event timing were synchronized via a Power1401 and recorded using ThorSync software.

#### Pharmacology

At the end of each experiment, a series of pharmacological agents were applied to identify sensory neuron subpopulations. Drugs targeted receptors highly expressed in the DRG and associated with specific neuronal subtypes based on single-cell sequencing data. Ligands included acetylcholine (1 mM), allyl isothiocyanate (AITC, a TRPA1 agonist;100 µM), bradykinin (1 µM), capsaicin (a TRPV1 agonist; 150 nM), cholecystokinin (CCK, a CCKBR agonist; 200 nM), chloroquine (a MrgprA3 agonist; 100 µM), histamine (500 µM), menthol (a TRPM8 agonist; 200 µM), oxotremorine (a non-selective muscarinic receptor agonist; 50 µM), oxytocin (1 µM), and prostaglandin E2 (PGE_2_; 50 nM). IL-1β (50 nM; R&D Systems, catalog #401-ML) was also applied to identify IL-1R1-expressing cells responsive to this ligand.

Following the pharmacology portion of the experiment, high potassium aCSF (30 mM) was perfused to activate all neurons and capture high-resolution images across all planes, confirming cell viability.

#### Analysis of calcium imaging data

Recording files were processed using Fiji (ImageJ) macros (available on GitHub: (https://github.com/cawarwick?tab=repositories). Suite2p (HHMI Janelia) handled image registration. Stabilized recordings were reviewed with another ImageJ macro to generate ROI summary images and averages for refinement. ROI detection was performed with Cellpose (v 2.2.3) refined manually in Fiji using functional responses. Only cells with stable XY position and no Z drift were analyzed. Following ROI selection, Fiji was used to extract raw mean fluorescence intensity. The ΔF/F was calculated in Python for each ROI using a rolling ball baseline: for each frame (Fi), the 30th percentile value of the surrounding 6.6 min of frames was used as baseline (Fb), and ΔF/F = (Fi - Fb)/Fb. Rolling ball normalization minimized photobleaching of GCaMP6s and minor changes in ROI location. ΔF/F traces were plotted over time and visually reviewed: ROIs showing Z drift, lack of K^+^ response, or cell death were excluded from the analysis. Neurons with basal spontaneous activity were counted but excluded from analyses of responsiveness to cutaneous stimuli or pharmacological agents.

### Statistical analysis

Statistical evaluations were performed using the Student’s t-test or one- or two-way ANOVA where appropriate. All statistical analyses were performed using Prism 10 (GraphPad Software Inc., v.10.6.0). A *p* value < 0.05 was considered as statistically significant. All data in graphs are expressed as means ± SEM.

## AUTHOR CONTRIBUTIONS

**Camille Illiano**: Conceptualization, Methodology, Formal Analysis, Investigation, Data Curation, Writing – Original Draft, Visualization. **Dominic Belanger:** Methodology, Formal Analysis, Investigation, Writing – Original Draft. **Harrison J. Stratton**: Methodology, Formal Analysis, Investigation, Writing – Original Draft. **Nicolas Vallières**: Conceptualization, Methodology, Formal Analysis, Visualization. **Nadia Fortin**: Methodology, Investigation.

**Martine Lessard**: Methodology, Investigation. **Benoit Mailhot**: Conceptualization, Methodology. **Louison Brochoire**: Methodology, Formal Analysis, Investigation. **Marine Christin**: Methodology, Formal Analysis, Investigation. **Yves De Koninck**: Resources. **Reza Sharif-Naeini**: Methodology, Formal Analysis, Investigation. **Sarah E. Ross**: Methodology, Formal Analysis, Investigation, Supervision. **Manon Defaye**: Methodology, Formal Analysis, Investigation. **Feng Wang**: Methodology, Formal Analysis, Investigation, Supervision. **Steve Lacroix**: Conceptualization, Methodology, Formal Analysis, Investigation, Writing – Original Draft, Writing – Review & Editing, Supervision, Project administration, Funding acquisition.

## ACKNOWLEDGEMENTS

This work was supported by the Canadian Institutes of Health Research (PJT-197909 to S.L., 2025-2030); MS Canada (Grant ID # 1416971 to S.L., 2025-2028); and the Wings for Life Spinal Cord Research Foundation (WFL-CA-18/24, Project # 316 to S.L., 2024-2027). Spatial transcriptomic studies were supported by a generous donation from the Club Richelieu de Sept-Îles. This work was also supported by the Fonds de recherche du Québec (FRQ) through a Research Centre grant to the Centre de recherche du CHU de Québec–Université Laval (Grant number 30641). During the preparation of this manuscript, the authors used Microsoft 365 Copilot to assist with grammar correction and readability improvements. All content was subsequently reviewed and edited by the authors, who take full responsibility for the final version of the published article.

## REFERENCES

Arbuckle, M.I., Komiyama, N.H., Delaney, A., Coba, M., Garry, E.M., Rosie, R., Allchorne, A.J., Forsyth, L.H., Bence, M., Carlisle, H.J., O’Dell, T.J., Mitchell, R., Fleetwood-Walker, S.M., Grant, S.G., 2010. The SH3 domain of postsynaptic density 95 mediates inflammatory pain through phosphatidylinositol-3-kinase recruitment. EMBO Rep 11, 473–478.

Arima, Y., Harada, M., Kamimura, D., Park, J.H., Kawano, F., Yull, F.E., Kawamoto, T., Iwakura, Y., Betz, U.A., Marquez, G., Blackwell, T.S., Ohira, Y., Hirano, T., Murakami, M., 2012. Regional neural activation defines a gateway for autoreactive T cells to cross the blood-brain barrier. Cell 148, 447–457.

Basbaum, A.I., Bautista, D.M., Scherrer, G., Julius, D., 2009. Cellular and molecular mechanisms of pain. Cell 139, 267–284.

Becht, E., McInnes, L., Healy, J., Dutertre, C.A., Kwok, I.W.H., Ng, L.G., Ginhoux, F., Newell, E.W., 2018. Dimensionality reduction for visualizing single-cell data using UMAP. Nat Biotechnol.

Bhuiyan, S.A., Nagi, S.S., Sankaranarayanan, I., Semizoglou, E., Usoskin, D., Yang, L., Yu, H., Arendt-Tranholm, A., Bertels, Z., Bhatia, P., Bouchatta, O., Boyer, K., Cervantes, A., Chalif, J., Chintalapudi, H., Cicalo, A., Copits, B., Cronin, C., Curatolo, M., Dong, X., Dougherty, P.M., Dourson, A., Funk, G., Gabriel, K., Griesemer, D.S., Guo, H., Gupta, P., Hofstetter, C., Horton, P., Hsieh, A., Inturi, N.N., Jain, A., Jayakar, S., Johnston, B., Kim, R., Krauter, D., Kupari, J., Lemen, J., Lesnak, J.B., Liu, W., Lopez, I., Lu, Y., MacMillan, H.J., Mazhar, K., Meriau, P., Moffitt, J.R., Moreno, M.M., Mwirigi, J.M., Naz, H., O’Brein, J., Payne, M., Del Rosario, J., Rosen, S.F., Shiers, S., Simpson, E., Slivicki, R., Stone, J.R., Tavares-Ferreira, D., Uhelski, M., Woolf, C.J., Xu, Q., Yi, J., Yousuf, M.S., Zhu, D., Cavalli, V., Zhao, G., Olausson, H., Ernfors, P., Gereau, R.W.t., Luo, W., Price, T.J., Renthal, W., Network, N.P.H.P., 2025. A Reference Atlas of the Human Dorsal Root Ganglion. bioRxiv.

Boilard, E., Nigrovic, P.A., Larabee, K., Watts, G.F., Coblyn, J.S., Weinblatt, M.E., Massarotti, E.M., Remold-O’Donnell, E., Farndale, R.W., Ware, J., Lee, D.M., 2010. Platelets amplify inflammation in arthritis via collagen-dependent microparticle production. Science 327, 580–583.

Bretheau, F., Castellanos-Molina, A., Belanger, D., Kusik, M., Mailhot, B., Boisvert, A., Vallieres, N., Lessard, M., Gunzer, M., Liu, X., Boilard, E., Quan, N., Lacroix, S., 2022. The alarmin interleukin-1alpha triggers secondary degeneration through reactive astrocytes and endothelium after spinal cord injury. Nat Commun 13, 5786.

Brown, E.V., Malik, A.F., Moese, E.R., McElroy, A.F., Lepore, A.C., 2022. Differential Activation of Pain Circuitry Neuron Populations in a Mouse Model of Spinal Cord Injury-Induced Neuropathic Pain. J Neurosci 42, 3271–3289.

Carlton, S.M., Du, J., Zhou, S., Coggeshall, R.E., 2001. Tonic control of peripheral cutaneous nociceptors by somatostatin receptors. J Neurosci 21, 4042–4049.

Caterina, M.J., Schumacher, M.A., Tominaga, M., Rosen, T.A., Levine, J.D., Julius, D., 1997. The capsaicin receptor: a heat-activated ion channel in the pain pathway. Nature 389, 816–824.

Chaplan, S.R., Bach, F.W., Pogrel, J.W., Chung, J.M., Yaksh, T.L., 1994. Quantitative assessment of tactile allodynia in the rat paw. J Neurosci Methods 53, 55–63.

Cook, A.D., Christensen, A.D., Tewari, D., McMahon, S.B., Hamilton, J.A., 2018. Immune Cytokines and Their Receptors in Inflammatory Pain. Trends Immunol 39, 240–255.

de Rivero Vaccari, J.P., Bastien, D., Yurcisin, G., Pineau, I., Dietrich, W.D., De Koninck, Y., Keane, R.W., Lacroix, S., 2012. P2X4 receptors influence inflammasome activation following spinal cord injury. J Neurosci 32, 3058–3066.

Defaye, M., Iftinca, M.C., Gadotti, V.M., Basso, L., Abdullah, N.S., Cumenal, M., Agosti, F., Hassan, A., Flynn, R., Martin, J., Soubeyre, V., Poulen, G., Lonjon, N., Vachiery-Lahaye, F., Bauchet, L., Mery, P.F., Bourinet, E., Zamponi, G.W., Altier, C., 2022. The neuronal tyrosine kinase receptor ligand ALKAL2 mediates persistent pain. J Clin Invest 132.

Economides, A.N., Carpenter, L.R., Rudge, J.S., Wong, V., Koehler-Stec, E.M., Hartnett, C., Pyles, E.A., Xu, X., Daly, T.J., Young, M.R., Fandl, J.P., Lee, F., Carver, S., McNay, J., Bailey, K., Ramakanth, S., Hutabarat, R., Huang, T.T., Radziejewski, C., Yancopoulos, G.D., Stahl, N., 2003. Cytokine traps: multi-component, high-affinity blockers of cytokine action. Nat Med 9, 47–52.

Fan, C.Y., McAllister, B.B., Stokes-Heck, S., Harding, E.K., Pereira de Vasconcelos, A., Mah, L.K., Lima, L.V., van den Hoogen, N.J., Rosen, S.F., Ham, B., Zhang, Z., Liu, H., Zemp, F.J., Burkhard, R., Geuking, M.B., Mahoney, D.J., Zamponi, G.W., Mogil, J.S., Ousman, S.S., Trang, T., 2025. Divergent sex-specific pannexin-1 mechanisms in microglia and T cells underlie neuropathic pain. Neuron 113, 896–911 e899.

Florio, S.K., Loh, C., Huang, S.M., Iwamaye, A.E., Kitto, K.F., Fowler, K.W., Treiberg, J.A., Hayflick, J.S., Walker, J.M., Fairbanks, C.A., Lai, Y., 2009. Disruption of nNOS-PSD95 protein-protein interaction inhibits acute thermal hyperalgesia and chronic mechanical allodynia in rodents. Br J Pharmacol 158, 494–506.

Fukuoka, H., Kawatani, M., Hisamitsu, T., Takeshige, C., 1994. Cutaneous hyperalgesia induced by peripheral injection of interleukin-1 beta in the rat. Brain Res 657, 133–140.

Gao, T., Dong, L., Qian, J., Ding, X., Zheng, Y., Wu, M., Meng, L., Jiao, Y., Gao, P., Luo, P., Zhang, G., Wu, C., Shi, X., Rong, W., 2021. G-Protein-Coupled Estrogen Receptor (GPER) in the Rostral Ventromedial Medulla Is Essential for Mobilizing Descending Inhibition of Itch. J Neurosci 41, 7727–7741.

Gaudet, A.D., Fonken, L.K., Ayala, M.T., Maier, S.F., Watkins, L.R., 2021. Aging and miR-155 in mice influence survival and neuropathic pain after spinal cord injury. Brain Behav Immun 97, 365–370.

Goldberg, D.S., McGee, S.J., 2011. Pain as a global public health priority. BMC Public Health 11, 770.

Grace, P.M., Hutchinson, M.R., Maier, S.F., Watkins, L.R., 2014. Pathological pain and the neuroimmune interface. Nat Rev Immunol 14, 217–231.

Hagen, K.M., Gordon, P., Frederick, A., Palmer, A.L., Edalat, P., Zonta, Y.R., Scott, L., Flancia, M., Reid, J.K., Joel, M., Ousman, S.S., 2024. CRYAB plays a role in terminating the presence of pro-inflammatory macrophages in the older, injured mouse peripheral nervous system. Neurobiol Aging 133, 1–15.

Hayashi, K., Lesnak, J.B., Plumb, A.N., Janowski, A.J., Smith, A.F., Hill, J.K., Sluka, K.A., 2024. Brain-derived neurotrophic factor contributes to activity-induced muscle pain in male but not female mice. Brain Behav Immun 120, 471–487.

Hu, X., Adebiyi, M.G., Luo, J., Sun, K., Le, T.T., Zhang, Y., Wu, H., Zhao, S., Karmouty-Quintana, H., Liu, H., Huang, A., Wen, Y.E., Zaika, O.L., Mamenko, M., Pochynyuk, O.M., Kellems, R.E., Eltzschig, H.K., Blackburn, M.R., Walters, E.T., Huang, D., Hu, H., Xia, Y., 2016. Sustained Elevated Adenosine via ADORA2B Promotes Chronic Pain through Neuro- immune Interaction. Cell Rep 16, 106–119.

Hu, X., Du, L., Liu, S., Lan, Z., Zang, K., Feng, J., Zhao, Y., Yang, X., Xie, Z., Wang, P.L., Ver Heul, A.M., Chen, L., Samineni, V.K., Wang, Y.Q., Lavine, K.J., Gereau, R.W.t., Wu, G.F., Hu, H., 2023. A TRPV4-dependent neuroimmune axis in the spinal cord promotes neuropathic pain. J Clin Invest 133.

Huang, J., Polgar, E., Solinski, H.J., Mishra, S.K., Tseng, P.Y., Iwagaki, N., Boyle, K.A., Dickie, A.C., Kriegbaum, M.C., Wildner, H., Zeilhofer, H.U., Watanabe, M., Riddell, J.S., Todd, A.J., Hoon, M.A., 2018. Circuit dissection of the role of somatostatin in itch and pain. Nat Neurosci 21, 707–716.

Huang, X., Deng, J., Xu, T., Xin, W., Zhang, Y., Ruan, X., 2021. Downregulation of metallothionein-2 contributes to oxaliplatin-induced neuropathic pain. J Neuroinflammation 18, 91.

Imamachi, N., Park, G.H., Lee, H., Anderson, D.J., Simon, M.I., Basbaum, A.I., Han, S.K., 2009. TRPV1-expressing primary afferents generate behavioral responses to pruritogens via multiple mechanisms. Proc Natl Acad Sci U S A 106, 11330–11335.

Inoue, A., Ikoma, K., Morioka, N., Kumagai, K., Hashimoto, T., Hide, I., Nakata, Y., 1999. Interleukin-1beta induces substance P release from primary afferent neurons through the cyclooxygenase-2 system. J Neurochem 73, 2206–2213.

James, G., Millecamps, M., Stone, L.S., Hodges, P.W., 2018. Dysregulation of the Inflammatory Mediators in the Multifidus Muscle After Spontaneous Intervertebral Disc Degeneration SPARC-null Mice is Ameliorated by Physical Activity. Spine (Phila Pa 1976) 43, E1184–E1194.

Jeon, S., Jha, M.K., Ock, J., Seo, J., Jin, M., Cho, H., Lee, W.H., Suk, K., 2013. Role of lipocalin-2-chemokine axis in the development of neuropathic pain following peripheral nerve injury. J Biol Chem 288, 24116–24127.

Ji, R.R., Chamessian, A., Zhang, Y.Q., 2016. Pain regulation by non-neuronal cells and inflammation. Science 354, 572–577.

Junger, H., Sorkin, L.S., 2000. Nociceptive and inflammatory effects of subcutaneous TNFalpha. Pain 85, 145–151.

Karchewski, L.A., Bloechlinger, S., Woolf, C.J., 2004. Axonal injury-dependent induction of the peripheral benzodiazepine receptor in small-diameter adult rat primary sensory neurons. Eur J Neurosci 20, 671–683.

Kawasaki, Y., Zhang, L., Cheng, J.K., Ji, R.R., 2008. Cytokine mechanisms of central sensitization: distinct and overlapping role of interleukin-1beta, interleukin-6, and tumor necrosis factor-alpha in regulating synaptic and neuronal activity in the superficial spinal cord. J Neurosci 28, 5189–5194.

Knight, B.E., Kozlowski, N., Havelin, J., King, T., Crocker, S.J., Young, E.E., Baumbauer, K.M., 2019. TIMP-1 Attenuates the Development of Inflammatory Pain Through MMP-Dependent and Receptor-Mediated Cell Signaling Mechanisms. Front Mol Neurosci 12, 220.

Kupari, J., Usoskin, D., Parisien, M., Lou, D., Hu, Y., Fatt, M., Lonnerberg, P., Spangberg, M., Eriksson, B., Barkas, N., Kharchenko, P.V., Lore, K., Khoury, S., Diatchenko, L., Ernfors, P., 2021. Single cell transcriptomics of primate sensory neurons identifies cell types associated with chronic pain. Nat Commun 12, 1510.

Kusik, M., Pare, A., Monteiro, F.D.G., Nadeau, S., Ferry, J., Pineau, I., Lessard, M., Fortin, N., Vallieres, N., Kerr, B., Lacroix, S., 2026. Inflammatory Monocyte-Derived Macrophages Drive Pain via Their Production of Nerve Growth Factor after Peripheral Nerve Injury in Mice. J Neurosci 46.

Kwilasz, A.J., Green Fulgham, S.M., Duran-Malle, J.C., Schrama, A.E.W., Mitten, E.H., Todd, L.S., Patel, H.P., Larson, T.A., Clements, M.A., Harris, K.M., Litwiler, S.T., Harvey, L.O., Jr., Maier, S.F., Chavez, R.A., Rice, K.C., Van Dam, A.M., Watkins, L.R., 2021. Toll-like receptor 2 and 4 antagonism for the treatment of experimental autoimmune encephalomyelitis (EAE)-related pain. Brain Behav Immun 93, 80–95.

Kwon, A., Jeon, S.M., Hwang, S.H., Kim, J.H., Cho, H.J., 2013. Expression and functional role of metallothioneins I and II in the spinal cord in inflammatory and neuropathic pain models. Brain Res 1523, 37–48.

Levesque, S.A., Pare, A., Mailhot, B., Bellver-Landete, V., Kebir, H., Lecuyer, M.A., Alvarez, J.I., Prat, A., de Rivero Vaccari, J.P., Keane, R.W., Lacroix, S., 2016. Myeloid cell transmigration across the CNS vasculature triggers IL-1beta-driven neuroinflammation during autoimmune encephalomyelitis in mice. J Exp Med 213, 929–949.

Li, X., Prudente, A.S., Prato, V., Guo, X., Hao, H., Jones, F., Figoli, S., Mullen, P., Wang, Y., Tonello, R., Lee, S.H., Shah, S., Maffei, B., Berta, T., Du, X., Gamper, N., 2024. Peripheral gating of mechanosensation by glial diazepam binding inhibitor. J Clin Invest 134.

Lim, E.F., Hoghooghi, V., Hagen, K.M., Kapoor, K., Frederick, A., Finlay, T.M., Ousman, S.S., 2021. Presence and activation of pro-inflammatory macrophages are associated with CRYAB expression in vitro and after peripheral nerve injury. J Neuroinflammation 18, 82.

Liu, X., Yamashita, T., Chen, Q., Belevych, N., McKim, D.B., Tarr, A.J., Coppola, V., Nath, N., Nemeth, D.P., Syed, Z.W., Sheridan, J.F., Godbout, J.P., Zuo, J., Quan, N., 2015. Interleukin 1 type 1 receptor restore: a genetic mouse model for studying interleukin 1 receptor-mediated effects in specific cell types. J Neurosci 35, 2860–2870.

Mailhot, B., Christin, M., Tessandier, N., Sotoudeh, C., Bretheau, F., Turmel, R., Pellerin, E., Wang, F., Bories, C., Joly-Beauparlant, C., De Koninck, Y., Droit, A., Cicchetti, F., Scherrer, G., Boilard, E., Sharif-Naeini, R., Lacroix, S., 2020. Neuronal interleukin-1 receptors mediate pain in chronic inflammatory diseases. J Exp Med 217.

Mapplebeck, J.C.S., Dalgarno, R., Tu, Y., Moriarty, O., Beggs, S., Kwok, C.H.T., Halievski, K., Assi, S., Mogil, J.S., Trang, T., Salter, M.W., 2018. Microglial P2X4R-evoked pain hypersensitivity is sexually dimorphic in rats. Pain 159, 1752–1763.

Merritt, C.R., Ong, G.T., Church, S.E., Barker, K., Danaher, P., Geiss, G., Hoang, M., Jung, J., Liang, Y., McKay-Fleisch, J., Nguyen, K., Norgaard, Z., Sorg, K., Sprague, I., Warren, C., Warren, S., Webster, P.J., Zhou, Z., Zollinger, D.R., Dunaway, D.L., Mills, G.B., Beechem, J.M., 2020. Multiplex digital spatial profiling of proteins and RNA in fixed tissue. Nat Biotechnol 38, 586–599.

Miclescu, A., Straatmann, A., Gkatziani, P., Butler, S., Karlsten, R., Gordh, T., 2019. Chronic neuropathic pain after traumatic peripheral nerve injuries in the upper extremity: prevalence, demographic and surgical determinants, impact on health and on pain medication. Scand J Pain 20, 95–108.

Mifflin, K.A., Kerr, B.J., 2017. Pain in autoimmune disorders. J Neurosci Res 95, 1282–1294.

Mogil, J.S., 2020. Qualitative sex differences in pain processing: emerging evidence of a biased literature. Nat Rev Neurosci 21, 353–365.

Mogil, J.S., Parisien, M., Esfahani, S.J., Diatchenko, L., 2024. Sex differences in mechanisms of pain hypersensitivity. Neurosci Biobehav Rev 163, 105749.

Moss, A., Ingram, R., Koch, S., Theodorou, A., Low, L., Baccei, M., Hathway, G.J., Costigan, M., Salton, S.R., Fitzgerald, M., 2008. Origins, actions and dynamic expression patterns of the neuropeptide VGF in rat peripheral and central sensory neurones following peripheral nerve injury. Mol Pain 4, 62.

Nadeau, S., Filali, M., Zhang, J., Kerr, B.J., Rivest, S., Soulet, D., Iwakura, Y., de Rivero Vaccari, J.P., Keane, R.W., Lacroix, S., 2011. Functional recovery after peripheral nerve injury is dependent on the pro-inflammatory cytokines IL-1{beta} and TNF: Implications for neuropathic pain. J Neurosci 31, 12533–12542.

Nampiaparampil, D.E., 2008. Prevalence of chronic pain after traumatic brain injury: a systematic review. JAMA 300, 711–719.

O’Connor, A.B., Schwid, S.R., Herrmann, D.N., Markman, J.D., Dworkin, R.H., 2008. Pain associated with multiple sclerosis: systematic review and proposed classification. Pain 137, 96–111.

Obreja, O., Schmelz, M., Poole, S., Kress, M., 2002. Interleukin-6 in combination with its soluble IL-6 receptor sensitises rat skin nociceptors to heat, in vivo. Pain 96, 57–62.

Okamoto, K., Ohashi, M., Ohno, K., Takeuchi, A., Matsuoka, E., Fujisato, K., Minami, T., Ito, S., Okuda-Ashitaka, E., 2016. Involvement of NIPSNAP1, a neuropeptide nocistatin- interacting protein, in inflammatory pain. Mol Pain 12.

Olechowski, C.J., Truong, J.J., Kerr, B.J., 2009. Neuropathic pain behaviours in a chronic-relapsing model of experimental autoimmune encephalomyelitis (EAE). Pain 141, 156–164.

Pineau, I., Lacroix, S., 2007. Proinflammatory cytokine synthesis in the injured mouse spinal cord: Multiphasic expression pattern and identification of the cell types involved. J Comp Neurol 500, 267–285.

Pinho-Ribeiro, F.A., Verri, W.A., Jr., Chiu, I.M., 2017. Nociceptor Sensory Neuron-Immune Interactions in Pain and Inflammation. Trends Immunol 38, 5–19.

Rahman, M.M., Hwang, S.M., Go, E.J., Kim, Y.H., Park, C.K., 2024. Irisin alleviates CFA-induced inflammatory pain by modulating macrophage polarization and spinal glial cell activation. Biomed Pharmacother 178, 117157.

Ren, K., Torres, R., 2009. Role of interleukin-1beta during pain and inflammation. Brain Res Rev 60, 57–64.

Safieh-Garabedian, B., Poole, S., Allchorne, A., Winter, J., Woolf, C.J., 1995. Contribution of interleukin-1 beta to the inflammation-induced increase in nerve growth factor levels and inflammatory hyperalgesia. Br J Pharmacol 115, 1265–1275.

Samad, T.A., Moore, K.A., Sapirstein, A., Billet, S., Allchorne, A., Poole, S., Bonventre, J.V., Woolf, C.J., 2001. Interleukin-1beta-mediated induction of Cox-2 in the CNS contributes to inflammatory pain hypersensitivity. Nature 410, 471–475.

Satija, R., Farrell, J.A., Gennert, D., Schier, A.F., Regev, A., 2015. Spatial reconstruction of single-cell gene expression data. Nat Biotechnol 33, 495–502.

Schafers, M., Sorkin, L., 2008. Effect of cytokines on neuronal excitability. Neurosci Lett 437, 188–193.

Sheahan, T.D., Warwick, C.A., Cui, A.Y., Baranger, D.A.A., Perry, V.J., Smith, K.M., Manalo, A.P., Nguyen, E.K., Koerber, H.R., Ross, S.E., 2024. Kappa opioids inhibit spinal output neurons to suppress itch. Sci Adv 10, eadp6038.

Silverman, W.R., de Rivero Vaccari, J.P., Locovei, S., Qiu, F., Carlsson, S.K., Scemes, E., Keane, R.W., Dahl, G., 2009. The pannexin 1 channel activates the inflammasome in neurons and astrocytes. J Biol Chem 284, 18143–18151.

Sonkoly, E., Muller, A., Lauerma, A.I., Pivarcsi, A., Soto, H., Kemeny, L., Alenius, H., Dieu-Nosjean, M.C., Meller, S., Rieker, J., Steinhoff, M., Hoffmann, T.K., Ruzicka, T., Zlotnik, A., Homey, B., 2006. IL-31: a new link between T cells and pruritus in atopic skin inflammation. J Allergy Clin Immunol 117, 411–417.

Sorge, R.E., LaCroix-Fralish, M.L., Tuttle, A.H., Sotocinal, S.G., Austin, J.S., Ritchie, J., Chanda, M.L., Graham, A.C., Topham, L., Beggs, S., Salter, M.W., Mogil, J.S., 2011. Spinal cord Toll-like receptor 4 mediates inflammatory and neuropathic hypersensitivity in male but not female mice. J Neurosci 31, 15450–15454.

Stemkowski, P.L., Bukhanova-Schulz, N., Baldwin, T., de Chaves, E.P., Smith, P.A., 2021. Are sensory neurons exquisitely sensitive to interleukin 1beta? J Neuroimmunol 354, 577529.

Tajerian, M., Alvarado, S., Millecamps, M., Dashwood, T., Anderson, K.M., Haglund, L., Ouellet, J., Szyf, M., Stone, L.S., 2011. DNA methylation of SPARC and chronic low back pain. Mol Pain 7, 65.

Tonello, R., Silveira Prudente, A., Hoon Lee, S., Faith Cohen, C., Xie, W., Paranjpe, A., Roh, J., Park, C.K., Chung, G., Strong, J.A., Zhang, J.M., Berta, T., 2023. Single-cell analysis of dorsal root ganglia reveals metalloproteinase signaling in satellite glial cells and pain. Brain Behav Immun 113, 401–414.

Tran, N.B., Lee, S.J., 2025. Metallothionein-3-mediated intracellular zinc mediates antioxidant and anti-inflammatory responses in the complete Freund’s adjuvant-induced inflammatory pain mouse model. Cell Death Discov 11, 45.

Usoskin, D., Furlan, A., Islam, S., Abdo, H., Lonnerberg, P., Lou, D., Hjerling-Leffler, J., Haeggstrom, J., Kharchenko, O., Kharchenko, P.V., Linnarsson, S., Ernfors, P., 2015. Unbiased classification of sensory neuron types by large-scale single-cell RNA sequencing. Nat Neurosci 18, 145–153.

Vicuna, L., Strochlic, D.E., Latremoliere, A., Bali, K.K., Simonetti, M., Husainie, D., Prokosch, S., Riva, P., Griffin, R.S., Njoo, C., Gehrig, S., Mall, M.A., Arnold, B., Devor, M., Woolf, C.J., Liberles, S.D., Costigan, M., Kuner, R., 2015. The serine protease inhibitor SerpinA3N attenuates neuropathic pain by inhibiting T cell-derived leukocyte elastase. Nat Med 21, 518–523.

Warwick, C., Salsovic, J., Hachisuka, J., Smith, K.M., Sheahan, T.D., Chen, H., Ibinson, J., Koerber, H.R., Ross, S.E., 2022. Cell type-specific calcium imaging of central sensitization in mouse dorsal horn. Nat Commun 13, 5199.

Yamanaka, H., Kobayashi, K., Okubo, M., Fukuoka, T., Noguchi, K., 2011. Increase of close homolog of cell adhesion molecule L1 in primary afferent by nerve injury and the contribution to neuropathic pain. J Comp Neurol 519, 1597–1615.

Zelenka, M., Schafers, M., Sommer, C., 2005. Intraneural injection of interleukin-1beta and tumor necrosis factor-alpha into rat sciatic nerve at physiological doses induces signs of neuropathic pain. Pain 116, 257–263.

Zhang, W., Xie, X., Xiong, X., Chen, F., 2025. HSPA1A Can Alleviate CFA-Induced Inflammatory Pain by Modulating Macrophages. Int J Mol Sci 26.

Zhang, Y., Gong, H., Wang, J.S., Li, M.N., Cao, D.L., Gu, J., Zhao, L.X., Zhang, X.D., Deng, Y.T., Dong, F.L., Gao, Y.J., Sun, W.X., Jiang, B.C., 2023. Nerve Injury-Induced gammaH2AX Reduction in Primary Sensory Neurons Is Involved in Neuropathic Pain Processing. Int J Mol Sci 24.

Zhao, W., Ma, L., Deng, D., Han, L., Xu, F., Zhang, T., Wang, Y., Huang, S., Ding, Y., Shu, S., Chen, X., 2025. BDNF-VGF Pathway Aggravates Incision Induced Acute Postoperative Pain via Upregulating the Neuroinflammation in Dorsal Root Ganglia. Mol Neurobiol 62, 169–183.

Zhou, Y., Zhou, B., Pache, L., Chang, M., Khodabakhshi, A.H., Tanaseichuk, O., Benner, C., Chanda, S.K., 2019. Metascape provides a biologist-oriented resource for the analysis of systems-level datasets. Nat Commun 10, 1523.

